# NtLLG4-mediated unconventional polar exocytosis of NtPPME1 coordinates cell wall rigidity and membrane dynamics to control pollen tube integrity

**DOI:** 10.1101/2025.02.24.639816

**Authors:** Xun Weng, Hao Wang, Yifeng Jiang, Ziheng Wang, Chuanhao Liu, Zhiheng Chen, Zhiyuan Yang, Jiayang Gao, Liwen Jiang, Lifeng Zhao, Jilei Huang, Hao Wang

**Author notes:** Corresponding author: Hao Wang, Ph.D., Professor.

## Abstract

Plant fertilization relies on controlled pollen tube growth that integrates membrane dynamics and cell wall expansion. We previously identified an unconventional exocytic pathway wherein Golgi-derived secretory vesicles (GDSVs) bypass the *trans*-Golgi network to deliver *Nicotiana tabacum* pectin methylesterase 1 (NtPPME1), thereby modulating cell wall rigidity. However, the mechanisms linking this patwhay with membrane dynamics and signaling remain elusive. Here, we used cryo-FIB-SEM and 3D tomography to identified GDSVs as a distinct vesicle population at the pollen tube tip. We further demonstrated that tobacco LORELEI-like-GPI-anchored protein 4 (NtLLG4), a key signaling molecule controlling membrane dynamics and integrity, functions as a receptor for NtPPME1, regulating its polar exocytosis *via* GDSVs to control cell wall stiffness. Furthermore, we identified trafficking signals which direct the unconventional exocytosis of NtPPME1 across intracellular organelles. Our findings reveal a crucial mechanism coupling cell wall rigidity with membrane signaling to control pollen tube growth and integrity during fertilization.

**Teaser:** We reveal a mechanism coupling cell wall rigidity with member signaling to control pollen tube growth and integrity during plant fertilization.

## Introduction

In angiosperms, sexual reproduction depends on the directional growth of pollen tubes to deliver two immotile male gametes to the female embryo sac for double fertilization (*1, 2*). Maintaining pollen tube integrity is critical during its rapid and polarized growth from the stigma surface towards the female gametophyte, ensuring that the pollen tube bursts at the appropriate time to release sperm cells upon arrival (*1–3*). An intricate signaling network including small peptides, receptor-like kinases (RLKs), actin cytoskeleton, Ca^2+^, pH and reactive oxygen species (ROS) in the pollen tube tip has been shown to maintain pollen tube integrity (*4–11*). Additionally, the rigidity of pollen tube cell wall must be spatiotemporally modulated to accommodate with membrane dynamics, allowing for active apical membrane expansion while maintaining the cylindrical shape of the pollen tube (*12–16*). However, the mechanisms coordinating membrane dynamics with cell wall rigidity still remain largely unknown. Apical vesicles, which carry newly synthesized proteins, lipid components, cell wall materials and extracellular signaling molecules accumulate in the tip region of growing pollen tubes to support and shape its growth (*9, 17–19*). Understanding the nature, dynamic behaviors and functions of these apical vesicles is of great importance to uncover the mechanisms underlying pollen tube growth and integrity.

Identification and studying the nature of apical vesicles in growing pollen tube tip is challenging due to their remarkably small size and vigorous behavior, which surpass the resolution limits of conventional light microscopy including confocal microscopy (*18, 20, 21*). Moreover, the lack of specific molecular markers for distinct types of vesicles in the tip region further complicates their differentiation (*18, 19, 22*). Previous studies using conventional transmission electron microscopy (TEM) have demonstrated that the apical vesicles enriched in the pollen tube tip are morphologically similar with a diameter of approximately 120 nm (*23–29*). However, Wang *et. al.*, have identified a distinct exocytic pathway in which Golgi-derived secretory vesicles (GDSVs) bypass the *trans*-Golgi network (TGN) to polarize the secretion of *Nicotiana tabacum* pectin methylesterase 1 (NtPPME1) at the apical surface of the growing pollen tube, regulating cell wall rigidity (*19*). This finding suggests that parallel exocytic pathway(s) may coexist and operate alongside the conventional TGN-derived exocytic pathway, facilitating the polar secretion of various cargoes to the pollen tube tip. Actually, GDSVs which are non-coated vesicles derived from Golgi apparatus with an approximate diameter of ∼40 nm, have been previously characterized based on their morphological features (*30*). Although NtPPME1 is known to be exocytosed *via* GDSVs into the apical apoplast to regulate pectic cell wall rigidity, two critical questions remain unanswered: i) Why have GDSVs not been observed and identified in the pollen tube tip region by conventional TEM imaging? While only 120 nm morphologically similar tip vesicles have been observed and identified (*23–29*); ii) Do NtPPME1-containing GDSVs actually traffic through the tip region of growing pollen tubes to reach the apical apoplast? This possibility remains speculative, as NtPPME1-GFP fluorescent puncta are not predominantly localized in the tip region but are instead enriched in distal regions further from the tip, as consistently demonstrated by several previous studies (*17, 31*). Emerging imaging technologies may provide new opportunities to elucidate the populations and dynamics of pollen tube tip vesicles. Cryo-focused ion beam scanning electron microscopy (cryo-FIB-SEM) combined with 3D tomography has emerged as a powerful tool for the visualization of ultrafine intracellular structures in three dimensions. This technique achieves high-resolution visualization by generating a series of continuous sections and SEM images while preserving intracellular structures in their near-native state through rapid freezing. It minimizes the potential risk of morphological alterations in endomembrane trafficking vesicles and organelles by obviating the need for subsequent sample substitution procedures with embedding resins (*23, 32–36*).

On the other hand, pollen tube integrity during fertilization has been shown to be regulated and sustained by the signaling peptides RALF4 and RALF19, which are derived from the pollen tube (*8, 9*). Upon the pollen tube’s arrival at the ovule, a related signaling peptide, RALF34, which secreted from female tissues, takes over and induces the rupture of the pollen tube (*8, 37*). The pollen tube perceives and response to these RALF peptide signals through an apical plasma membrane (PM)-localized receptor-coreceptor complex composed of the RLKs ANXUR1/2 (ANX1/2), Buddha’s Paper Seal 1/2 (BUPS1/2) and pollen tube-specific GPI-APs LORELEI-LIKE GPI-ANCHORED PROTEIN 2/3 (LLG2/3) (*6, 8, 38*). The polar localization of this protein complex at the pollen tube apex is mediated by transport and deposition through tip vesicles (*6, 38*). This membrane-associated RALF-ANX-BUPS-LLG signaling complex regulates downstream Ca²⁺ gradients and ROS levels to ensure pollen tube integrity and polar growth (*6, 8, 9, 38, 39*). Loss-of-function mutations or reduced expression of RALF4/19, LLG2/3, ANX1/2, or BUPS1/2 consistently result in early pollen tube burst prior to reaching the ovules (*6, 8, 38–40*). Meanwhile, the rigidity of the pollen tube cell wall must be spatially regulated to coordinate with membrane dynamics during growth. The apical cell wall must remain sufficiently plastic to allow rapid membrane expansion at the tip or facilitate pollen tube burst when releasing sperm cells, while the cell wall in the shank region should be stiff enough to withstand the turgor pressure and maintain the cylindrical shape of the pollen tube (*12–15, 41, 42*). However, it remains largely uncovered how the underlying mechanisms coordinate cell wall rigidity with membrane signaling to maintain pollen tube integrity.

In this study, we employed cryo-FIB-SEM and 3D tomography to identify GDSVs as a distinct type of apical vesicle accumulating at the pollen tube tip for the first time. Additionally, we systematically dissected the specific trafficking signals that guide NtPPME1 through the GDSV-mediated unconventional polar exocytosis. We further demonstrated that NtLLG4, a member of the RALF-ANX-BUPS-LLG signaling complex involved in PM dynamics and integrity, serves as a receptor for NtPPME1, regulating its polar secretion *via* GDSVs to modulate cell wall rigidity at the pollen tube tip. Our findings reveal a vital mechanism that synchronizes cell wall rigidity with membrane signaling to preserve pollen tube integrity during fertilization, providing significant insights into the biological functions of GDSV-mediated unconventional exocytosis in polar cell growth and morphogenesis during plant fertilization.

## Results

### Morphological identification and 3D tomography of GDSVs in NtPPME1-mediated unconventional polar exocytosis at the pollen tube tip

To elucidate the mechanisms underlying NtPPME1-mediated unconventional polar exocytosis, it is essential to first morphologically identify GDSVs within the pollen tube tip. The tip region is densely populated with numerous tiny vesicles, which have been shown in previous studies to be morphologically similar (*18, 19, 23, 29, 43, 44*). These vesicles are believed to be involved in the dynamic processes of exocytosis, endocytosis and recycling, which occur concurrently at the pollen tube tip (*18, 19, 45*). However, it remains unclear how a single vesicle type can mediate these three distinct intracellular trafficking pathways, or whether multiple types of apical vesicles co-exist in the pollen tube tip.

To accurately locate and morphologically identify the vesicles in the pollen tube tip, we employed cryo-FIB-SEM, an advanced approach distinct from conventional TEM sample preparation and imaging methods. This method allowed us to capture serial sections of *Nicotiana tobacum* pollen tube tip and generates high-resolution 3D tomography for the first time. The general sample preparation and workflow of cryo-FIB-SEM and 3D tomography were illustrated in Fig. 1A-D. Mature pollens were germinated *in vitro* for two hours and subsequently vitrified on a TEM grid by plunge-freezing (Fig. 1A). The grid was then mounted into the FIB-SEM cryo-shuttle and the vitrified pollen tubes were identified using the SEM prior to FIB milling. A germinated pollen tube on the grid was selected (Fig. 1B), with the tip region was specifically targeted for FIB sectioning (Fig. 1C). A series of continued sections, each 30 nm in thickness, was obtained from the pollen tube tip through FIB milling and subsequently imaged using SEM (Fig. 1D). Finally, the ultrafine 3D structures within the tip region were reconstructed into a 3D tomography, as illustrated in Fig. 1D.

**Fig. 1.**
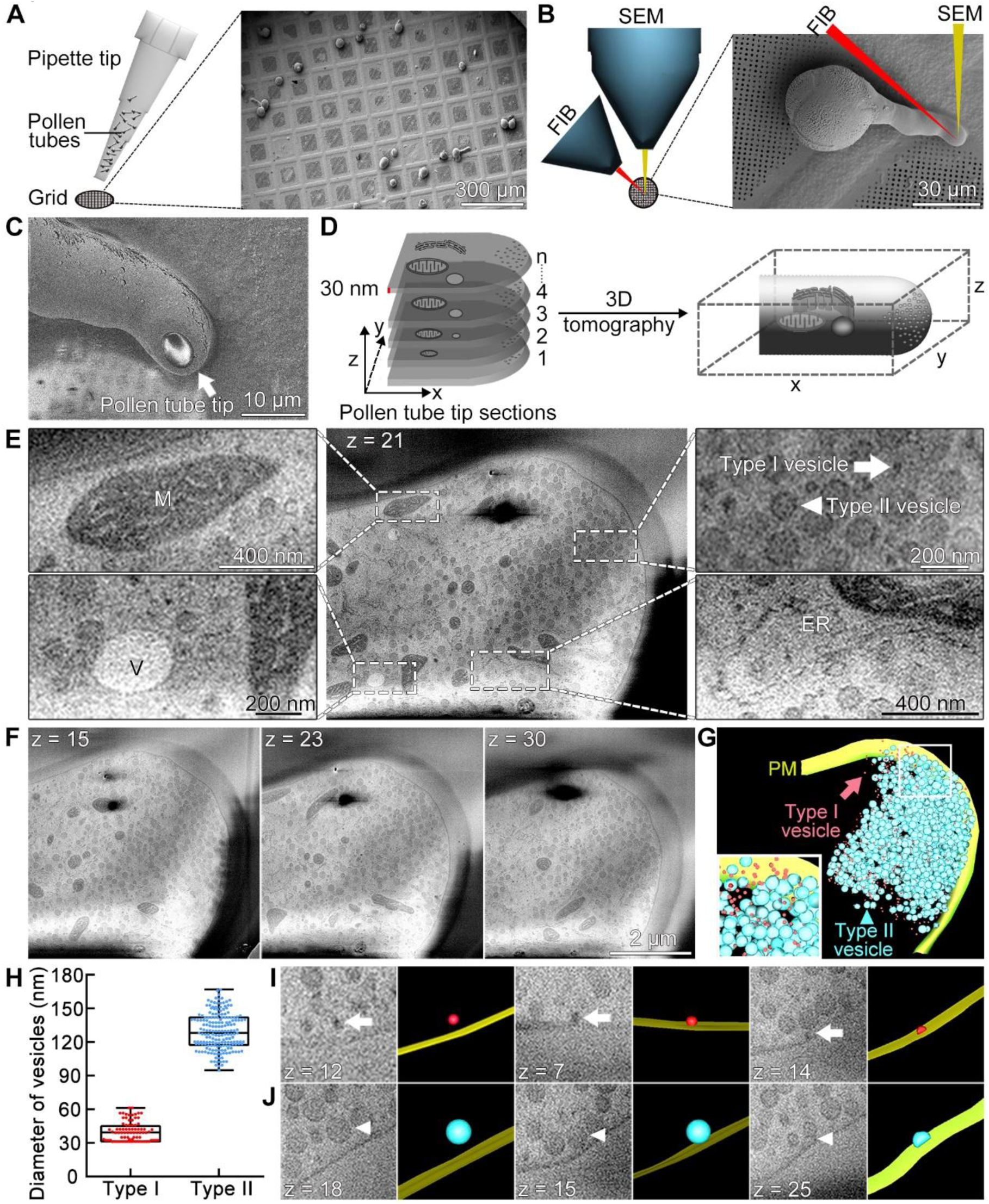
Cryo-FIB-SEM morphological identification of GDSVs in the tobacco pollen tube tip. (**A**) Schematic illustration of the preparation of germinated pollen tubes on a gold TEM grid, with an enlarged view showing the tubes on the grid. (**B**) Simplified schematic of the principle of FIB milling and SEM imaging, with an enlarged view highlighting the focus on the germinated pollen tube tip. (**C**) Representative image of a germinated pollen tube tip after FIB milling. (**D**) Schematic demonstration of the consecutive 30 nm-thick sections of pollen tube tip obtained by cryo-FIB-SEM. Subsequent 3D tomography of pollen tube was generated through image stacking and computational modeling. (**E**) Representative SEM image of a cryo-FIB slide from the pollen tube showing well-preserved organelles, including mitochondria, vacuoles, the ER, and two morphologically distinct apical vesicles named Type I and Type II. (**F**) SEM images from different cryo-FIB slices of the pollen tube tip. (**G**) 3D tomography of Type I and Type II apical vesicles within the pollen tube tip. (**H**) Statistical analysis of the diameters of Type I and Type II apical vesicles. (**I**) **and** (**J**) Representative images and 3D tomography illustrating the dynamics of Type I and Type II vesicles including their proximity to the PM, contact with the PM and interact with the PM. Arrows in (**E**), (**G**) and (**I**) indicate Type I vesicle. Arrowheads in (**E**), (**G**) and (**J**) indicates Type II vesicle. M, mitochondrion. V, vacuole. ER, endoplasmic reticulum.

The intracellular structures in the pollen tube tip region including organelles such as mitochondria, small vacuoles, and the endoplasmic reticulum (ER) in the subapical region, are well-preserved and morphologically clear (Fig. 1E). Notably, we identified two distinct types of non-coated vesicles in the apical region of the pollen tube tip (Fig. 1E and movie S1). Type I vesicles exhibit high electron density and have an average diameter of 40 nm (n=88) (Fig. 1E-H and movie S1). The morphology of the Type I vesicles is in consistence with the GDSVs which have been identified as mediators of unconventional exocytosis of NtPPME1 bypassing the TGN (*19*). In contrast, Type II vesicles are characterized by a relatively lower electron density and a larger average diameter of 130 nm (n=185) (Fig. 1E-H and movie S1). The morphology of Type II vesicles aligns with previous observations of apical vesicles in pollen tubes (*23, 29, 43*). To further investigate these vesicles, we generated 3D tomography of the apical vesicles that are densely accumulated at the pollen tube tip (Fig. 1F and G), using serial continued sections of SEM images. This allowed us to visualize the spatial distribution and morphologies of both Type I and Type II vesicles (Fig. 1G). In addition, we observed that both types of apical vesicles are capable of contacting and interacting with the PM (Fig. 1I and J and movie S1). These findings demonstrate that GDSVs are localized in the growing pollen tube tip and have the capacity to interact with and fuse to the PM, supporting their role in NtPPME1-mediated unconventional polar exocytosis.

### Dissecting domain-specific trafficking signal determinants of NtPPME1 during unconventional polar exocytosis

To gain deeper insights into the intracellular trafficking signals that regulate the unconventional polar exocytic process of NtPPME1, we systematically examined the functions of its different domains through a series of truncations, deletions, and point mutations. Using InterPro (https://www.ebi.ac.uk/interpro/), we predicted the primary structure of NtPPME1, which includes a 23-amino acid signal peptide (SP), following followed by a pro-region, a random coil (RC) region, and a C-terminal PME domain as illustrated in Fig. 2A. NtPPME1 is classified as the type II PME subfamily, characterized by a unique pre-pro-protein structural configuration that encompasses both the PME inhibitor (PMEI) domain and the PME catalytic domain (*46, 47*). The pro-region of NtPPME1 has been demonstrated to act as a PMEI, effectively inhibits the enzymatic activity of PME during its intracellular trafficking. Upon exocytosis of NtPPME1 into the apoplast, its enzymatic activity is activated through the cleavage at the RC region, which separates the pro-region from the PME region (*48, 49*). Moreover, to further explore the spatial organization of these domains, we used AlphaFold3 for structural modeling (fig. S1 and Fig. 2B). The predicted structures of the pro-region and PME domain were determined with high confidence, as reflected by predicted local distance difference test (pLDDT) scores exceeding 85 (high confidence: 70 < pLDDT < 100) (fig. S1). The pro-region primarily consists of multiple α-helices, while the PME domain is largely composed of β-sheets (Fig. 2B). The conserved structural features revealed by these high-confidence models highlight the evolutionary importance of these domains for proper function, supporting the need for accurate structural predictions.

**Fig. 2.**
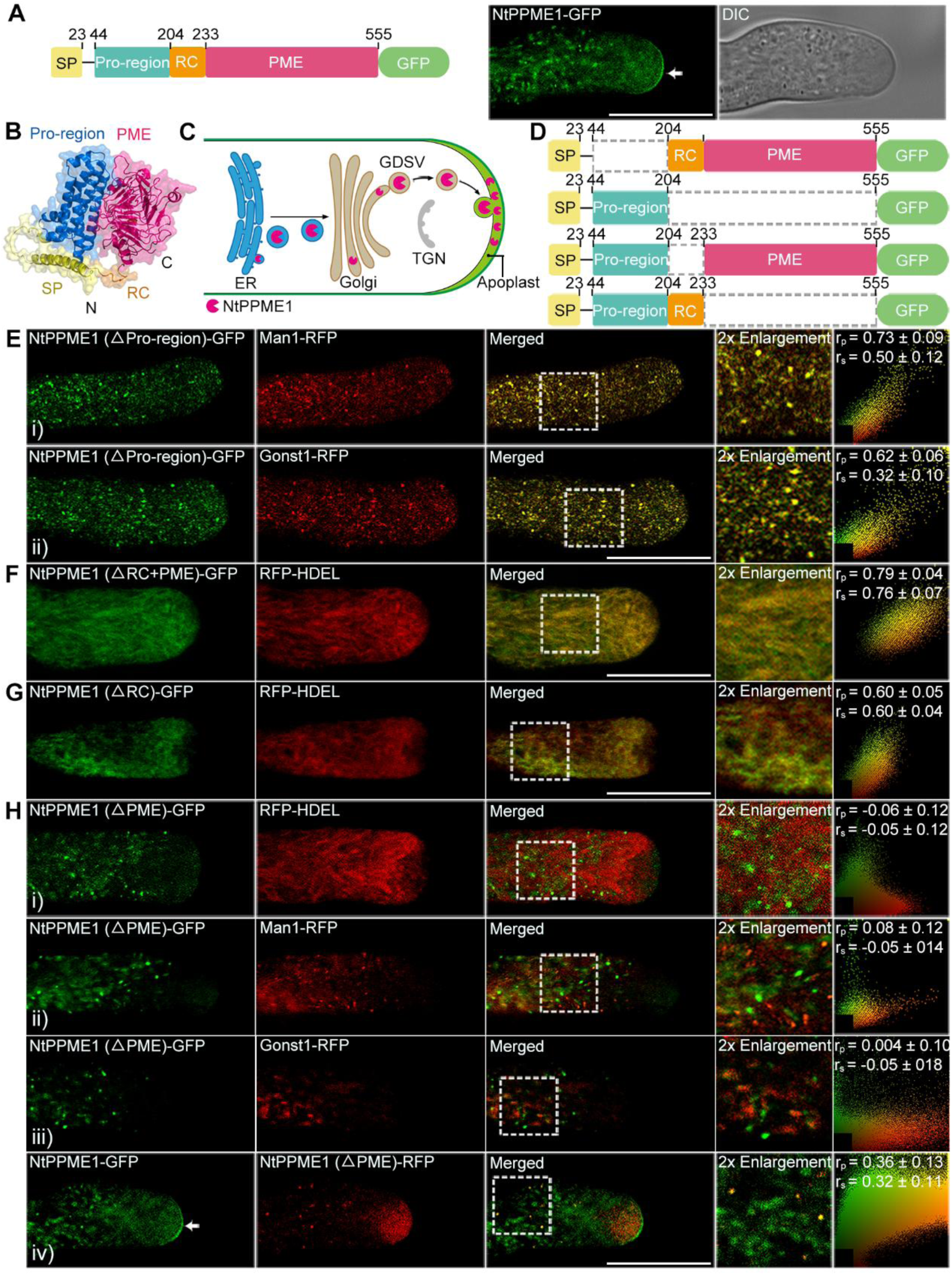
Dissection of the domain-specific trafficking signals of NtPPME1 during polar unconventional exocytosis. (**A**) Schematic illustration of the primary structure of NtPPME1 protein (left) and representative images of a tobacco growing pollen tube expressing NtPPME1-GFP. (**B**) Predicted 3D structure of NtPPME1 by AlphaFold3. (**C**) Schematic representation of the unconventional exocytosis of NtPPME1 in the growing pollen tube. (**D**) Schematic of truncated NtPPME1 constructs fused with GFP. (**E**) Representative images of tobacco growing pollen tubes co-expressing NtPPME1 (ΔPro-region)-GFP with **i)** Man1-RFP or **ii)** Gonst1-RFP. (**F**) Representative image of tobacco growing pollen tubes co-expressing NtPPME1 (ΔRC+PME)-GFP and RFP-HDEL. (**G**) Representative image of tobacco growing pollen tubes co-expressing NtPPME1 (ΔRC)-GFP and RFP-HDEL. (**H**) Representative images of tobacco growing pollen tubes co-expressing NtPPME1 (ΔPME)-GFP/RFP with **i)** RFP-HDEL, **ii)** Man1-RFP, **iii)** Gonst1-RFP or **iv)** NtPPME1-GFP. Colocalization ratios were calculated using Pearson correlation coefficients or Spearman’s rank correlation coefficients with ImageJ. The r values in the range of −1 to 1, with 0 indicating no discernable correlation and +1 or −1 indicating strong positive or negative correlations, respectively. White arrows in (**A**) and (**H**) indicate apical localization of NtPPME1-GFP at the growing pollen tube tip. SP, signal peptide. Pro-region, pro-region domain. RC, random coil. PME, pectin methylesterase domain. N, N-terminus. C, C-terminus. ER, endoplasmic reticulum. GDSV, Golgi-derived secretory vesicle. TGN, *trans*-Golgi network. Scale bar in (**A**), (**E**), (**F**), (**G**) and (**H**): 15 μm.

To assess whether the pro-region, RC region, and PME domain of NtPPME1 contain trafficking signals mediating its unconventional polar exocytic pathway, we generated chimeric proteins by fusing truncated versions of NtPPME1 with GFP, then transiently expressed them in growing tobacco pollen tubes (Fig. 2C-H and movie S3-S10). Deletion of the pro-region, RC region, or PME domain completely abolished the polar localization of NtPPME1 at the pollen tube tip compared to the full-length protein (Fig. 2E-H). Additionally, these deletions showed distinct intracellular localizations and distributions. To precisely determine the subcellular localizations of these truncated variants, we compared them with various organelle markers (Fig. 2E-H and movies S3-S10). Our results found that NtPPME1 (ΔPro-region)-GFP colocalized with both Man1-RFP and Gonst1-RFP, which serve as markers for the *cis*-Golgi and *trans*-Golgi, respectively (Fig. 2E and movies S3 and S4). Deletion of the RC region and PME domain caused NtPPME1 (ΔRC+PME)-GFP to be retained in the ER, as evidenced by its colocalization with RFP-HDEL, an ER marker (Fig. 2F and movie S5). Additionally, deletion of the RC region also caused NtPPME1 (ΔRC)-GFP to be retained in the ER (Fig. 2G and movie S6). Although deletion of the PME domain resulted in the loss of apical localization at the pollen tube tip, NtPPME1 (ΔPME)-GFP remained localized to GDSVs due to its colocalization with NtPPME1 intracellular puncta, while it remained separate from ER and Golgi markers (Fig. 2H and movies S7-S10). This suggests that the PME domain may contain a polar exocytic signal essential for its targeting to the pollen tube tip. In addition, the subcellular localizations of NtPPME1 (ΔRC+PME)-GFP, NtPPME1 (ΔPME)-GFP and NtPPME1 (ΔRC)-GFP suggest that the RC region exclusively contains the export signal necessary for directing the secretion of NtPPME1 from the ER to the Golgi apparatus. Similarly, the localizations of NtPPME1 (ΔPro-region)-GFP and NtPPME1 (ΔPME)-GFP indicate that the pro-region may harbor a Golgi export signal responsible for trafficking from the Golgi to GDSVs. Together, our results reveal that each domain of NtPPME1 contains distinct organelle trafficking signals that collectively drive its unconventional polar exocytosis, bypassing the TGN.

### Identification of ER-to-Golgi and Golgi export signals in NtPPME1 regulating unconventional polar exocytosis in pollen tubes

To further elucidate the specific intracellular trafficking signals that regulate the unconventional polar exocytosis of NtPPME1 in growing pollen tubes, we systematically generated various deletions and point mutations in the C-terminus of the RC, as the RC region is known to contain the ER-to-Golgi trafficking signal (Fig. 2F-H and 3A). Our data revealed that progressive deletions of the last six to sixteen amino acids at the C-terminus of the RC region resulted in increased co-localization with the ER (Fig. 3A panel i and ii, fig. S2A and movies S11 and S12). This suggests that the last 16 amino acids (FVEASNRKLLQISNAK) within the RC region harbor the ER export signal essential for NtPPME1 trafficking to the Golgi. Notably, the 16 amino acids contain dihydrophobic (LL) and FV motifs, previously identified as ER export signals in mammals and yeast (*50–52*). Mutations in the FV_218, 219_ and LL_225, 226_ motifs, either individually or in combination, resulted in partial or complete co-localization with the ER marker (Fig. 3A panel iii, fig. S2B and movie S13). These findings confirm that the FV_218, 219_ and LL_225, 226_ within the C-terminal region of the RC domain are crucial for mediating the trafficking of NtPPME1 from the ER to the Golgi apparatus (Fig. 3B).

**Fig. 3.**
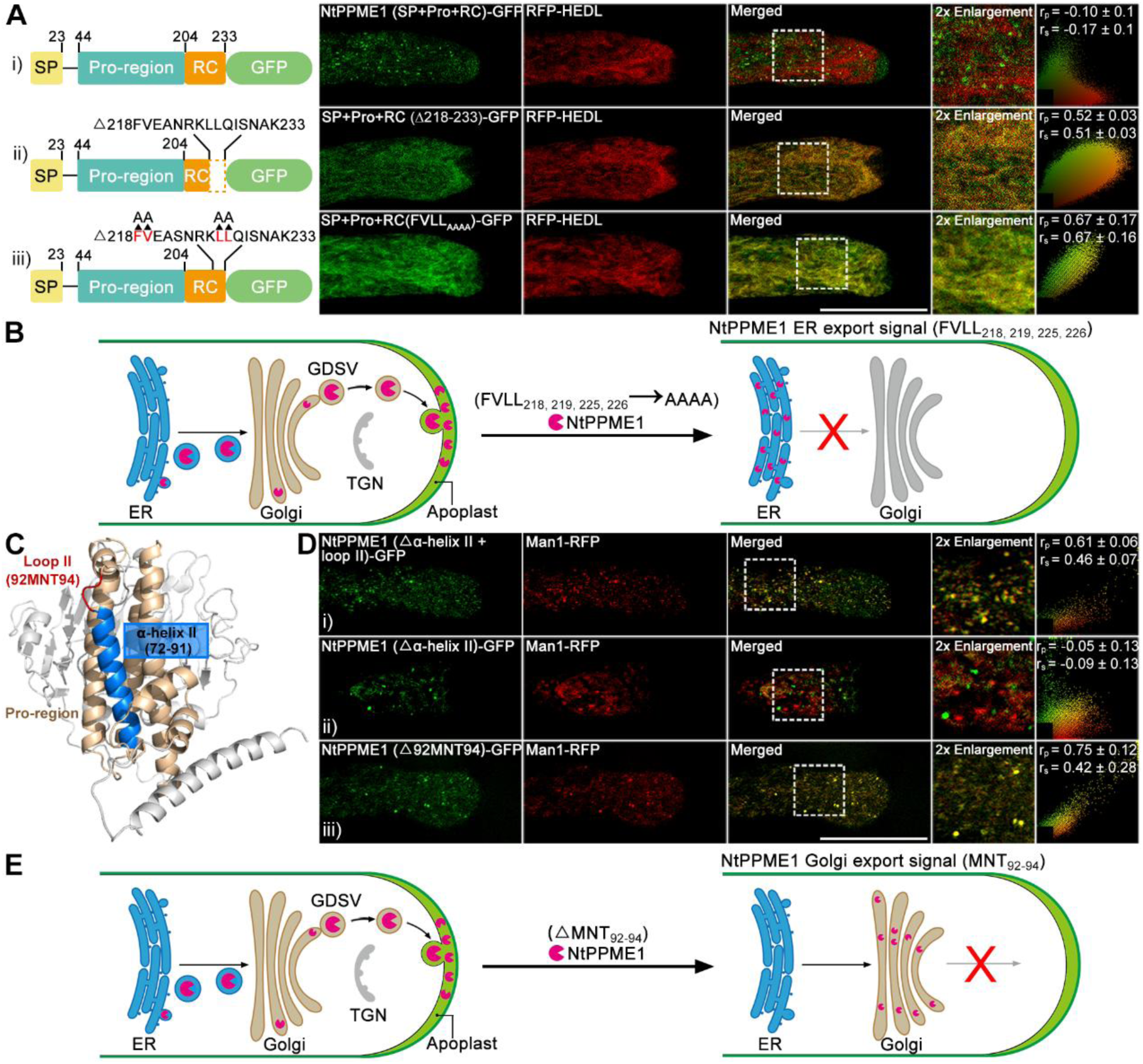
Identification of specific trafficking signals mediating NtPPME1 secretion from the ER to GDSV. (**A**) Representative image of tobacco growing pollen tubes co-expressing of the ER marker RFP-HDEL with **i)** NtPPME1 (SP+Pro+RC)-GFP, **ii)** SP+Pro+RC(Δ218-233)-GFP and **iii)** SP+Pro+RC(FVLL_AAAA_)-GFP. (**B**) Schematic identification of the ER export determinant trafficking signal FVLL_218, 219, 225, 226_ in the RC region of NtPPME1 during unconventional polar exocytosis. (**C**) Schematic highlight the structures of α-helix II and the loop II region connecting it with α-helix III in the Pro-region of NtPPME1. (**D**) Representative images of tobacco pollen tubes co-expressing of the Golgi marker Man1-RFP with **i)** NtPPME1 (Δα-helix II + loop II)-GFP, **ii)** NtPPME1 (Δα-helix II)-GFP or **iii)** NtPPME1 (Δ92MNT94)-GFP. (**E**) Schematic identification of the Golgi export signal MNT_92-94_ in the pro-region of NtPPME1 during unconventional polar exocytosis. Colocalization ratios were calculated using either Pearson correlation coefficients or as Spearman’s rank correlation coefficients with imageJ. The r values in the range of −1 to 1, with 0 indicate no discernable correlation and +1 or −1 indicates strong positive or negative correlations, respectively. SP, signal peptide. Pro-region, pro-region domain. RC, random coil. ER, endoplasmic reticulum. GDSV, Golgi-derived secretory vesicles. TGN, *trans*-Golgi network. Scale bars in (**A**) and (**D**): 15 μm.

Next, we investigated the Golgi export signal within the pro-region of NtPPME1. Given that truncation of the pro-region led to the retention of NtPPME1 (ΔPro-region)-GFP within the Golgi apparatus, this observation suggests that the pro-region contains the Golgi export signal (Fig. 2E). Using AlphaFold3, we predicted that the pro-region is composed of five α-helices (I-V) interconnected by short loops (fig. S3A). To pinpoint the Golgi export signal, we generated GFP-tagged NtPPME1 variants with sequential deletions of each α-helix and their adjacent loops (Δ45-71, Δ72-94, Δ95-138, Δ139-169, and Δ170-204) (Fig. 3D panel i, fig. S3B and movie S14). Deletion of the second α-helix and the loop II (72–94) resulted in Golgi retention, indicating that this region contains the Golgi export signal (Fig. 3D, panel i and movie S14). In contrast, deletion of the other α-helixes and their adjacent loops led to punctate distribution, separating from the Golgi apparatus (fig. S3B). Further truncations of α-helix II (spanning amino acids 72-91) and the loop (MNT_92-94_) linking α-helix II and III (Fig. 3D panel ii and iii, and movie S15 and S16) revealed that the MNT_92-94_ loop functions as the Golgi export signal (Fig. 3D panel iii and movie S16). Thus, we identified MNT_92-94_ within the pro-region of NtPPME1 as the Golgi export signal responsible for mediating the exit of NtPPME1 from the Golgi apparatus (Fig. 3E).

### Identification of key trafficking signals and a glycosylation site regulating NtPPME1 polar exocytosis from GDSVs to the apoplast

To elucidate the signals that regulate the unconventional polar exocytosis of NtPPME1 from GDSVs to the apoplast, we generated a series of deletions at the C-terminus of the PME domain (Fig. 4A), as the PME domain is believed to contain the GDSV-to-apoplast trafficking signal for NtPPME1 (Fig. 2A and H). Deletion of the last four amino acids (MMKV) at the C-terminus of the PME domain did not affect the apical localization of the NtPPME1 mutant at the pollen tube tip, which remained consistent with the full-length protein (Fig. 4A panel i and movie S17). However, deletion of the last seven amino acids (EAGMMKV) completely abolished this polar localization (Fig. 4A panel ii and movie S18). This suggests that the amino acids EAG_549-551_ at the C-terminus is essential for polar exocytosis to the apoplast. To further validate this finding, we generated GFP-tagged mutants with single, double, and triple amino acid changes in the EAG_549-551_ motif (fig. S4A and B, Fig. 4A panel iii, and movie S19). Both mutation and deletion of the EAG_549-551_ motif disrupted the polar localization of NtPPME1 at the pollen tube tip (Fig. 4A panel iii and fig. S4C). In contrast, single and double amino acid mutations had no effect (fig. S4A and B). In conclusion, the EAG_549-551_ motif in the C-terminal region of the PME domain is critical for mediating the trafficking of NtPPME1 from GDSVs to the apoplast (Fig. 4C).

**Fig. 4.**
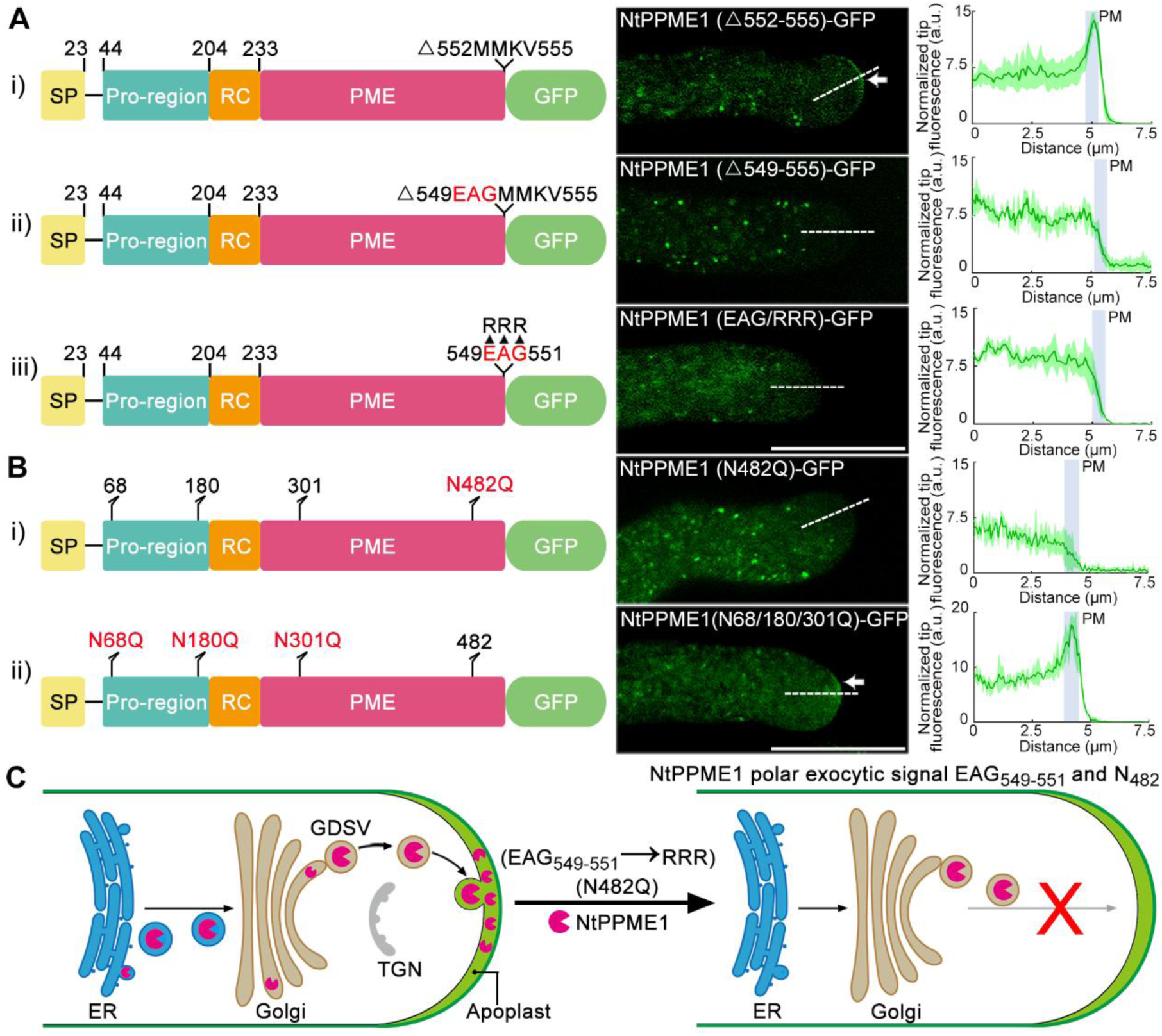
Identification of the exocytic determinant signals controlling NtPPME1 unconventional polar exocytosis from GDSVs to apoplast. (**A**) Chimeric GFP fusions with progressive C-terminal deletions or key amino acids mutation of the PME region in NtPPME1 and representative images of tobacco pollen tubes expressing these deletion plasmids. (**B**) Chimeric GFP fusions with various mutations at the four N-glycosylation sites of NtPPME1, and representative images of pollen tubes expressing these mutated constructs. White arrows indicate the apical localization of expressed protein at the growing pollen tube tip. Measurements of apical fluorescence signal intensity across the pollen tube tip region as indicated by the white dashed lines are normalized and plotted into graphs (n ≥ 3). (**C**) Schematic identification of the polar exocytic signal of EAG_549-551_ and the N-glycosylation site N_482_ in the C-terminal of PME region of NtPPME1 controls unconventional polar exocytosis. SP, signal peptide. Pro-region, pro-region domain. RC, random coil. PME, pectin methylesterase domain. PM, plasma membrane. ER, endoplasmic reticulum. GDSV, Golgi-derived secretory vesicles. TGN, *trans*-Golgi network. Scale bars in (**A**) and (**B**): 15 μm.

In addition, glycosylation is a common post-translational modification that influences protein localization and ensures proper folding (*53, 54*). To determine whether glycosylation plays a role in NtPPME1 trafficking, we used NetNGlyc (https://services.healthtech.dtu.dk/services/NetNGlyc-1.0/) to predict the N-linked glycosylation sites in NtPPME1 (fig. S5A). Four putative sites were identified at N68, N180, N301, and N482 (fig. S5A). We then generated GFP-tagged point mutations at these positions (N68Q, N180Q, N301Q, and N482Q) revealed that only the N482Q mutation disrupted apical localization (Fig. 4B, panel i and movie S20). The other mutations, either alone or in combination, had no effect (Fig. 4B panel ii, fig. S5B and movie S21). Furthermore, the N482Q mutation caused NtPPME1 to be retained in GDSVs, as indicated by its colocalization with NtPPME1 intracellular puncta, while it remained separate from Golgi apparatus and TGN markers (fig. S5C). These findings demonstrate that the N-glycosylation site at N482 crucial for mediating NtPPME1 trafficking from GDSVs to the apoplast (Fig. 4C).

### NtLLG4 acts as a key receptor in NtPPME1-mediated unconventional polar exocytosis and targeting

Due to the limited understanding of GDSV-mediated unconventional polar exocytosis involved in pollen tube polar growth and cell plate formation during cytokinesis as revealed in previous studies, NtPPME1 remains the only characterized cargo protein associated with this pathway (*19, 31, 44*). In addition, it is highly speculated that the specific intracellular sorting and packaging of soluble NtPPME1 which undergoes through the unconventional polar exocytosis pathway is likely to be a receptor-mediated active process. Therefore, to test this hypothesis and investigate which receptor protein mediates the targeting of soluble cargo NtPPME1 to the pollen tube tip, we isolated total proteins from transgenic tobacco pollen tubes expressing NtPPME1-GFP and performed immunoprecipitation (IP) using GFP-Trap, followed by high-performance liquid chromatography-tandem mass spectrometry (HPLC-MS/MS) analysis. The identified proteins were classified into four functional categories based on Eukaryotic Orthologous Groups (KOG) annotation, with the largest group associated with cellular processes and signaling (fig. S6A and table S1). Given that NtPPME1 is a secreted soluble cargo protein containing a SP, we focused our analysis on 165 potential receptor proteins that contain either an SP or a transmembrane domain. Among these, we identified a glycosylphosphatidylinositol-anchored protein (GPI-AP), LOC107817512, which functions as a chaperone and receptor in maintaining pollen tube integrity in Arabidopsis (fig. S6B) (*6, 38*). This receptor was designated as *Nicotiana tabacum* LORELEI-like GPI-AP 4 (NtLLG4).

To further investigate the GPI-AP family in *Nicotiana tabacum*, we used the NtLLG4 sequence as a query in BLAST searches, identifying nine members of the NtLLG family (fig. S7A). Comparative analysis with Arabidopsis LLGs (AtLLGs) revealed that three of the nine NtLLGs belong to the AtLORELEI and AtLLG1 clade, which are known to regulate root growth in Arabidopsis (*38, 55*). The remaining six NtLLGs cluster within the AtLLG2/3 clade, which is involved in regulating pollen tube integrity (fig. S7A) (*6, 38, 55*). Sequence alignment confirmed a high degree of conservation among the NtLLG family proteins (fig. S7B). Furthermore, gene expression profiling using the NCBI database demonstrated that *NtLLG4* and *NtLLG5* are highly expressed in *Nicotiana tabacum* flowers across various developmental stages, including young, mature, and senescent stages (fig. S7C), consistent with the gene expression patterns of *AtLLG2/3* in Arabidopsis (*38*).

To further elucidate the spatial interaction between NtPPME1 and NtLLG4, we performed protein structural modeling of both proteins using AlphaFold3 (Fig. 5A-C). The structural models were predicted with high confidence, as evidenced by a predicted template modeling (pTM) score exceeding 0.9 (pTM > 0.8 represents confident high-quality predictions) (fig. S8). Notably, the protruding surface region of NtLLG4 (highlighted with the black dashed cycle) interacts with the concave region of NtPPME1 (highlighted with the white dashed cycle) (Fig. 5C). This concave region of NtPPME1 is formed by integration of the pro-region (dark blue) and the PME domain (magenta), suggesting that NtLLG4 interacts with both domains of NtPPME1 (Fig. 5C). Furthermore, co-localization analysis revealed that the punctate signals of NtPPME1-GFP predominantly co-localized with NtLLG4-RFP (Fig. 5D). To explore the interaction in more detail, we conducted a co-IP assay using anti-GFP and anti-Myc antibodies by transiently co-expressing NtPPME1-GFP and NtLLG4-Myc in *Nicotiana benthamiana* leaves. It confirmed that NtPPME1 interacts with NtLLG4 (Fig. 5E). Moreover, fluorescence resonance energy transfer (FRET) analysis and luciferase complementation assay (LCA) further validated this interaction in tobacco BY-2 protoplasts and in plants (Fig. 5F and G). Besides, a yeast two-hybrid (Y2H) assay also confirmed that NtPPME1 interacts with NtLLG4 in the absence of their N-terminal SP (Fig. 5H).

**Fig. 5.**
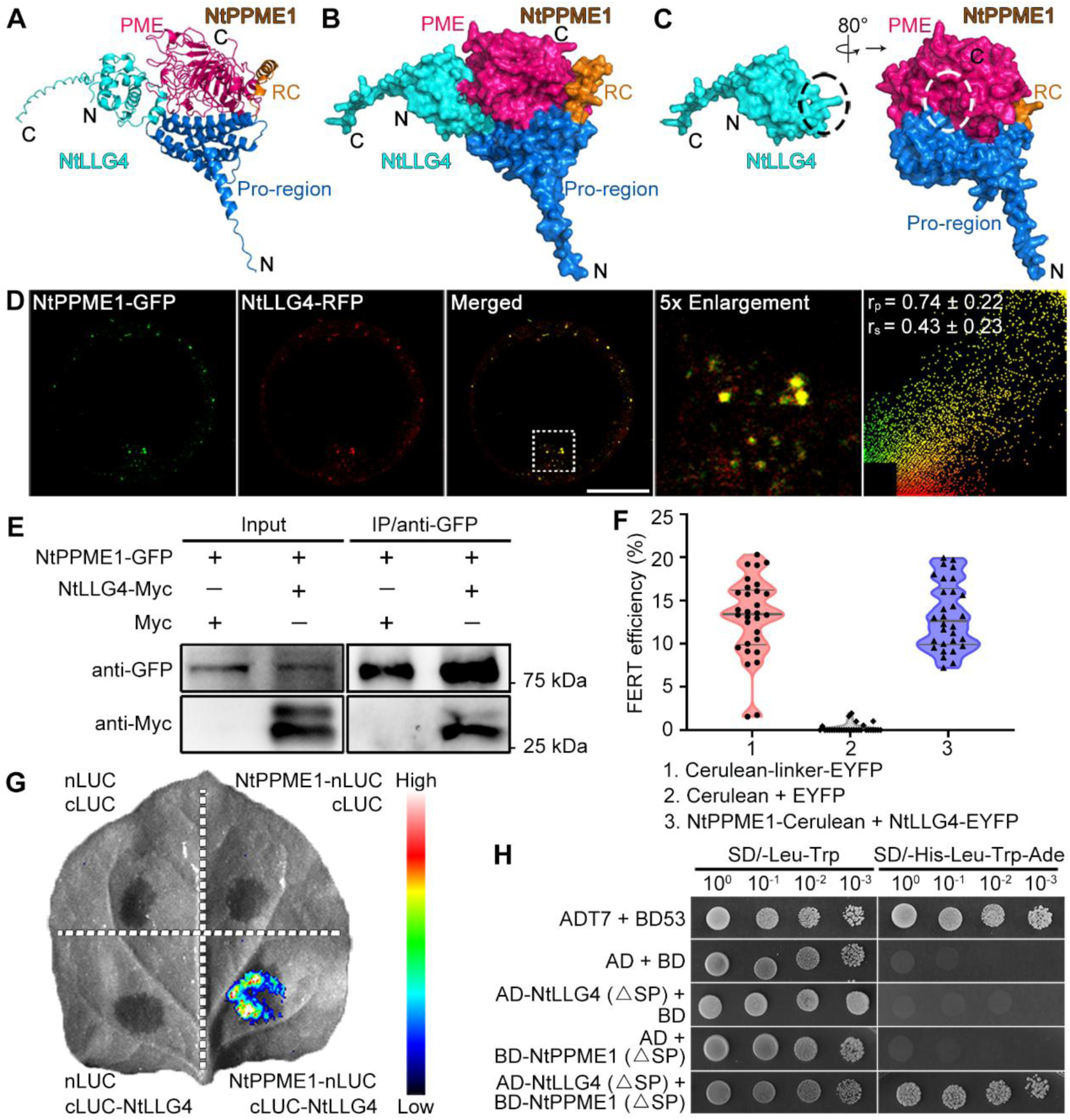
Identification of NtLLG4 acting as a receptor for NtPPME1 during unconventional polar exocytosis. (**A**) and (**B**) Structural and surface predictions of the NtLLG4-NtPPME1 protein-protein interaction using AlphaFold3. (**C**) The protein-protein interaction interfaces between NtLLG4 and NtPPME1, as predicted by AlphaFold3, are highlighted by black and white dashed circles, respectively. (**D**) Representative images of tobacco BY-2 protoplast co-expressing NtPPME1-GFP and NtLLG4-RFP. Scale bar: 10 μm. Colocalization ratios are calculated using either Pearson correlation coefficients or as Spearman’s rank correlation coefficients with image J. The generated r values in the range of −1 to 1, with 0 indicating no discernable correlation and +1 or −1 indicating strong positive or negative correlations, respectively. (**E**) Co-IP analysis demonstrating the interaction between NtPPME1 and NtLLG4. (**F**) FRET analysis of the protein interaction between NtPPME1 and NtLLG4 in BY-2 protoplasts. (**G**) LCA to assess the interaction between NtPPME1 and NtLLG4. The *Nicotiana benthamiana* leaves were divided into four sections and infiltrated with different combinations of expression plasmids as indicated. (**H**) Y2H analysis exploring the roles of NtPPME1 in the interaction between NtPPME1 and NtLLG4. SP, signal peptide. Pro-region, pro-region domain. RC, random coil. PME, pectin methylesterase domain. N, N-terminus. C, C-terminus.

To dissect the interaction sites, we generated NtPPME1 truncations targeting either the pro-region or the PME of NtPPME1. Co-IP assays, Y2H analysis, and LCA all demonstrated that NtLLG4 interacts with both the pro-region and PME domain of NtPPME1 (Fig. 6A-C). These findings suggest that NtLLG4 acts as a specific receptor that binds NtPPME1 during its unconventional polar exocytosis prior to its release into the apoplast of growing pollen tubes. To further investigate the specific interaction sites between NtPPME1 and NtLLG4, we conducted protein-protein interaction surface analyses of the NtPPME1-NtLLG4 complex using PDBePISA (https://www.ebi.ac.uk/pdbe/pisa/) (Fig. 6D). The analysis revealed that NtLLG4 establishes electrostatic interactions with the surface of NtPPME1 (Fig. 6D). Specifically, the R41 residue of NtLLG4 fits into an acidic pocket on NtPPME1, engaging residues E165, Q342, Q364, D365, Y368, R454, and N482, which distributed across the pro-region and PME domains (Fig. 6D i and ii). Additionally, R41 of NtLLG4 forms hydrogen bonds with D365 of NtPPME1 (Fig. 6D ii). Furthermore, N51 of NtLLG4 forms a hydrogen bond with E173 of NtPPME1 *via* its nitrogen and oxygen atoms, respectively (Fig. 6D iii and iv). Mutational analysis using Y2H and LCA showed that mutations in NtPPME1 (EEQQDYRN_AAAAAAAQ_) disrupted the interaction between NtPPME1 and NtLLG4 (Fig. 6E and F). These results confirm that the interaction of NtLLG4 to both the pro-region and PME domain of NtPPME1 is primarily mediated through a network of hydrogen bonds and electrostatic interactions.

**Fig. 6.**
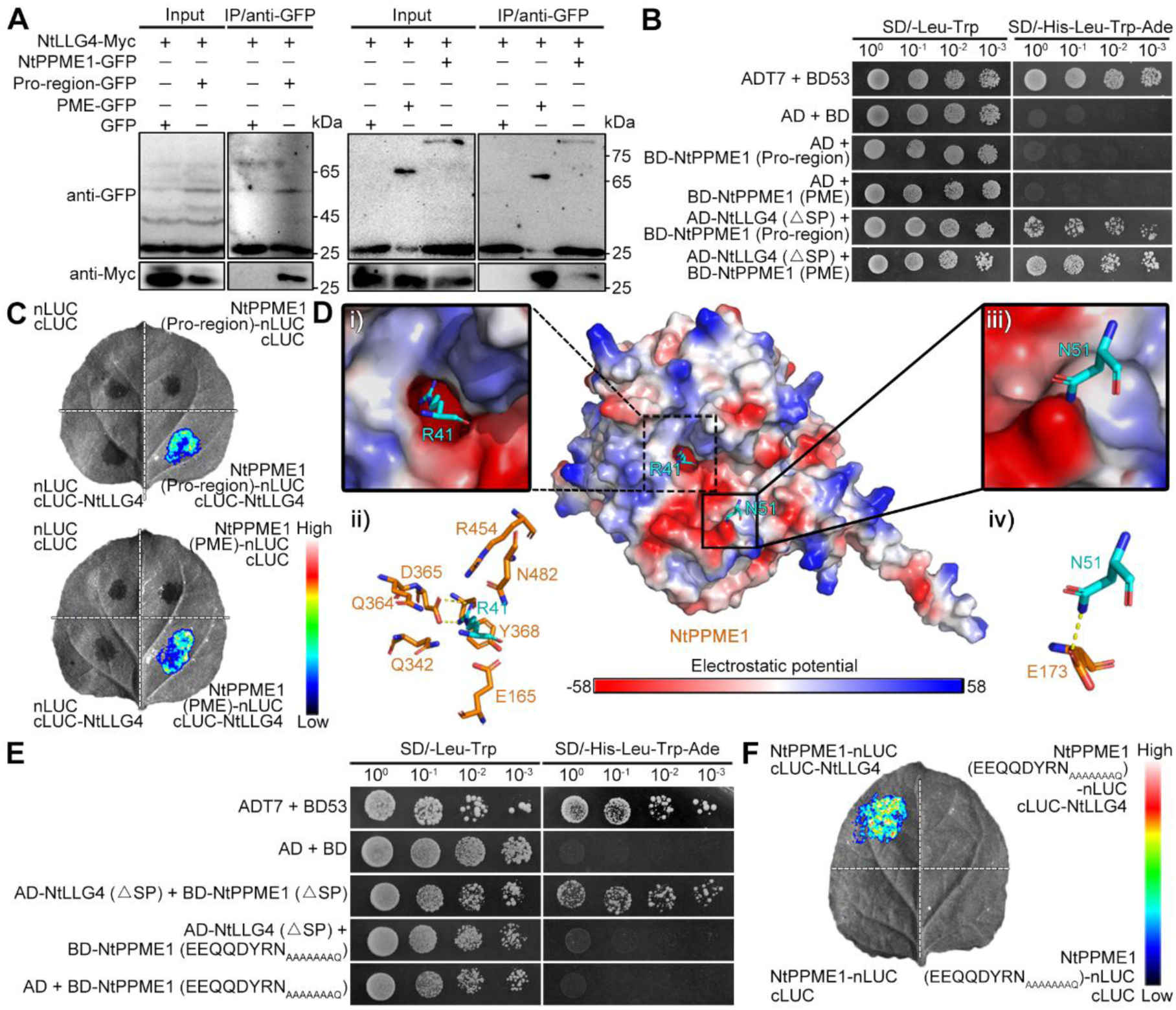
Reveal the structural interaction sites between NtPPME1 and NtLLG4. (**A**) Co-IP analysis investigating the roles of the pro-region and PME domains in the interaction between NtPPME1 and NtLLG4. (**B**) Y2H analysis exploring the roles of the pro-region and PME domains in the interaction between NtPPME1 and NtLLG4. (**C**) LCA to assess the roles of the pro-region and PME domains in the interaction between NtPPME1 and NtLLG4. (**D**) Predicted specific protein binding sites between NtPPME1 and NtLLG4. NtPPME1 amino acids are highlighted in orange, while NtLLG4 amino acids are shown in blue. The positively and negatively charged surfaces of NtPPME1 are represented in blue and red, respectively. (**E**) Y2H analysis examining the interaction between NtPPME1 mutant (mutations at E165, E173, Q342, Q364, D365, Y368, R454, N482_AAAAAAAQ_) and NtLLG4. (**F**) LCA accessing the interaction between NtPPME1 mutant (mutations at E165, E173, Q342, Q364, D365, Y368, R454, N482_AAAAAAAQ_) and NtLLG4. SP, signal peptide. Pro-region, pro-region domain. RC, random coil. PME, pectin methylesterase domain. N, N-terminus. C, C-terminus.

### NtLLG4 regulates NtPPME1 polar exocytosis and pectic cell wall construction to sustain pollen tube integrity

Given the interaction between NtLLG4 and NtPPME1, we next sought to determine the biological functions of NtLLG4 in determining the polar localization of NtPPME1 at the pollen tube tip, and its effects on pollen tube germination and growth. Co-expression of NtPPME1-GFP and NtLLG4-RFP in tobacco pollen tubes revealed that both proteins co-localized as punctate structures (Fig. 7A and movie S22). Nevertheless, it is noteworthy that the fluorescent signal of NtPPME1-GFP was absent at the pollen tube apex when NtLLG4 was overexpressed (Fig. 7A and B). To further explore the function of NtLLG in pollen tube growth, we used RNAi to silence the NtLLG family, avoiding potentially functional redundancy among family members. Upon knockdown of the *NtLLG* gene family, the tip localization of NtPPME1-RFP was also abolished (Fig. 7C and movie S23). Either the overexpression of NtLLG4 or *NtLLG-RNAi* can result in the disruption of apical localization of NtPPME1 at the pollen tube tip. It indicates that accurate amount of NtLLG4 receptors is required to facilitate the unconventional polar exocytosis of NtPPME1. It is further supported by the previous functional studies of LLG protein in maintaining pollen tube integrity and controlling cell polarity, which LLG level in growing pollen tubes need to be precisely regulated (*6, 38*). Additionally, approximately 40% of pollen tubes co-expressing NtPPME1-RFP and *NtLLG-RNAi* exhibited abnormally multiple pollen germination sits and tube growth (Fig. 7D and E). It indicates critical roles of NtLLG in guarding a single site for pollen germination and proper tube growth.

**Fig. 7.**
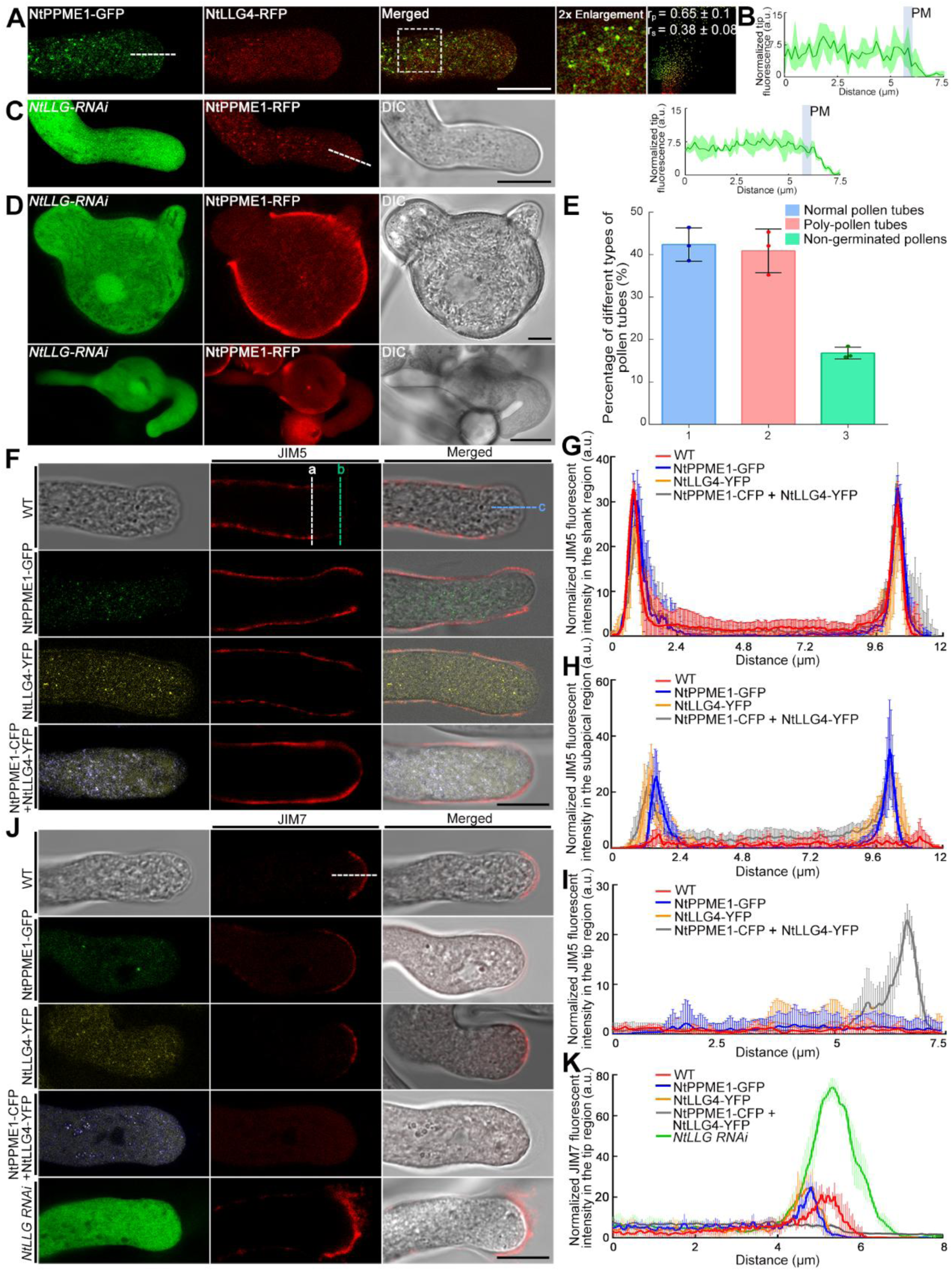
Coordination between NtPPME1 and NtLLG4 in regulating pollen tube cell wall rigidity. (**A**) Representative images of pollen tubes co-expressing NtPPME1-GFP and NtLLG4-RFP. Scale bar: 15 μm. (**B**) The intensity of NtPPME1-GFP at the apical tip were measured along the white dashed line indicated in (**A**) and plotted on a graph (n = 3). (**C**) Representative images of pollen tubes co-expressing *NtLLG-RNAi* and NtPPME1-RFP. The intensity of NtPPME1-RFP at the apical tip were measured along the white dashed line (left) and plotted on a graph (n = 3). Scale bar: 15 μm. (**D**) Representative images of co-expressing *NtLLG4-RNAi* and NtPPME1-RFP caused germination and growth of poly-pollen tubes from a single pollen grain. Scale bar: 25 μm (upper panel), 8 μm (lower panel). (**E**) Statistical analysis of the percentage of different phenotypes observed in (**D**). The results represent the means ± SEs (n ≥ 125). (**F**) Immunofluorescent labeling of demethylesterified pectin distribution in pollen tubes. Three different lines (a, b, and c) were drawn across the shank, subapical, and tip regions of the pollen tube, respectively, for signal measurement and analysis. Scale bar: 15 μm. (**G**) to (**I**) Measurement of JIM5 fluorescence in the shank, subapical, and tip regions of the pollen tube, as indicated by the dashed lines (a, b, c) in (**F**), and plotted on a graph (n ≥ 4), respectively. (**J**) Immunolabeling of methylesterified pectin distribution in pollen tubes. Scale bar: 15 μm. (**K**) Measurement of JIM7 fluorescence across the pollen tube tip region, as indicated by the white dashed line in (**J**), and plotted on a graph (n ≥ 5).

To further reveal the functional coordination between NtLLG4 and NtPPME1 in pollen tube growth, we modulated the expression levels of *NtPPME1* and/or *NtLLG4* and analyzed the resulting changes in cell wall rigidity using immunofluorescent labeling with JIM5 and JIM7 antibodies. The pollen tube tip cell wall is primarily composed of highly methylesterified pectin, recognized by the JIM7 antibody, which provides the flexibility necessary for the rapid expansion of the pollen tube membrane (*28, 56, 57*). As the pollen tube elongates, the esterified pectin is progressively demethylesterified by PME, recognized by the JIM5 antibody, leading to a stiffer cell wall that supports the cylindrical structure of the tube (*12, 44*). Compared to WT pollen tubes, those expressing either NtPPME1-GFP or NtLLG4-YFP showed increased cell wall stiffness in the subapical region as indicated by an extended JIM5 signal (Fig. 7F-H). The fluorescent signal intensity of JIM5 was quantitatively assessed across the shank and subapical regions of pollen tubes, and plotted on a graph (n ≥ 4) (Fig. 7G and H). Moreover, in pollen tubes co-expressing NtPPME1-CFP and NtLLG4-YFP, JIM5 fluorescence extended into both the subapical and apical regions, indicating increased rigidity throughout the tip (Fig. 7F and I). JIM5 fluorescence was quantitatively measured across the apical region of pollen tubes, and plotted on a graph (n = 5) (Fig. 7I). Consistent with the JIM5 labeling results, the JIM7 fluorescence signal at the pollen tube apex was reduced in tubes expressing either NtPPME1-GFP or NtLLG4-YFP, and was completely absent in tubes co-expressing both proteins (Fig. 7J and K). It suggests that co-expression of NtPPME1 and NtLLG4 enhances pectin demethylesterification at the pollen tube tip, contributing to increased cell wall rigidity. Conversely, a significantly increased JIM7 fluorescent signal was observed at the apex of *NtLLG-RNAi* pollen tubes (Fig. 7J and K), indicating that NtLLG4 is essential for the proper regulation of pectin methylesterification. Quantitative analysis of JIM7 fluorescence in the apical region was plotted (n ≥ 5) (Fig. 7K). These findings indicate that NtLLG4 acts as a receptor for NtPPME1, collaboratively regulating its polar exocytosis and targeting at the pollen tube apex. Their interaction modulates the rigidity of the pollen tube cell wall, allowing for the spatiotemporal coordination with membrane signaling required to maintain pollen tube integrity during plant fertilization.

Collectively, we propose a schematic model to illustrate the cooperative mechanism of cell wall rigidity and membrane signaling in controlling pollen tube integrity during plant fertilization (Fig. 8). Each domain of NtPPME1 harbors specific trafficking signals that mediate its unconventional polar exocytosis within the growing pollen tube. NtLLG4 serves as a receptor to interact with NtPPME1, ensuring its targeted exocytic secretion to the apoplast. Upon release into the apoplast, NtPPME1 undergoes proteolytic cleavage, separating the pro-region from the PME domain which then catalyzes pectin demethylesterification to modulate cell wall rigidity. This process ensures the proper morphology and structural integrity of the pollen tube. Furthermore, NtLLG4 is a crucial component of the RALF-LLG-ANX-BUPS signaling complex on the PM, which regulates pollen tube integrity and timely rupture (*6, 38*). Thus, our study uncovers a coordinated mechanism between cell wall rigidity and membrane signaling that governs pollen tube integrity during plant fertilization.

**Fig. 8.**
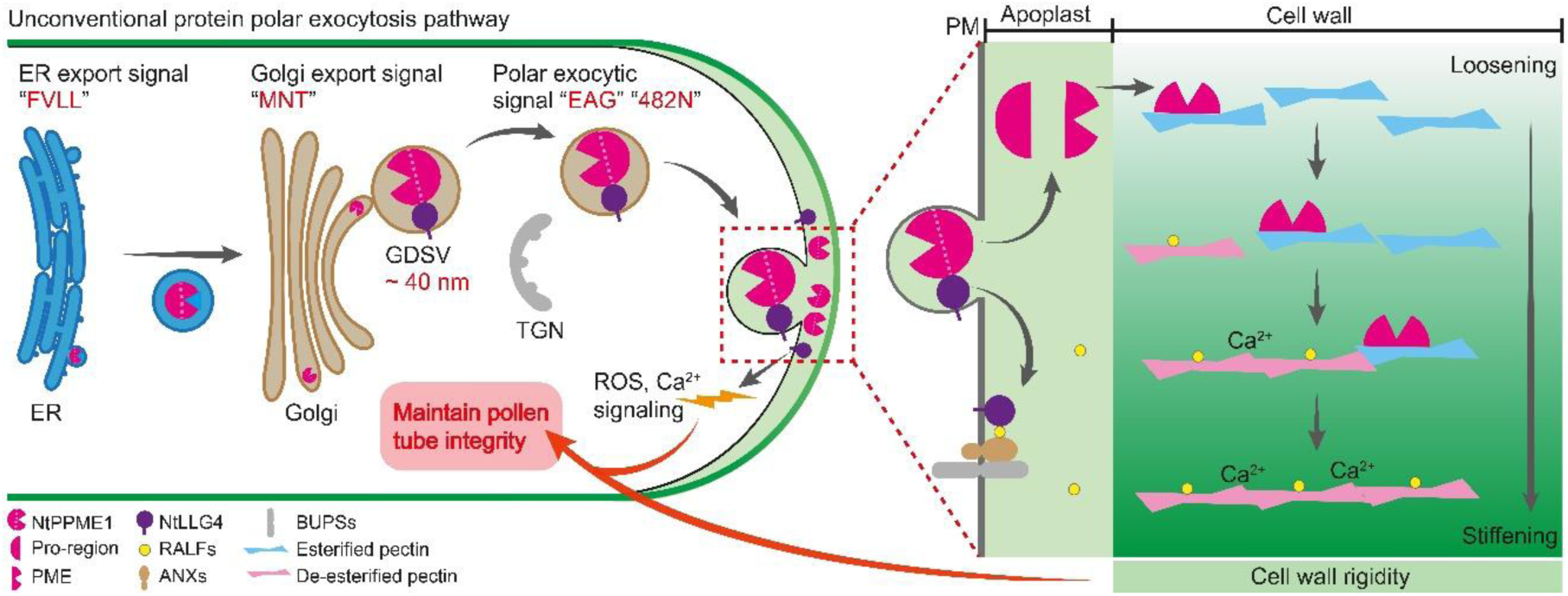
Schematic hypothetical model of the regulatory mechanism underlying NtPPME1 unconventional polar exocytosis and its role in maintaining pollen tube integrity. This schematic illustrates a proposed model for the regulatory mechanisms by which NtPPME1 mediated unconventional polar exocytosis and contributes to the maintenance of pollen tube integrity through cell wall rigidity. Each domain of NtPPME1 contains specific organelle trafficking signals that enable its polar exocytosis in the growing pollen tube. During this process, NtLLG4 interacts with NtPPME1, directing its targeted trafficking to the pollen tube apex. Once released into the apoplast, NtPPME1 undergoes proteolytic cleavage, separating the PME domain from the pro-region. The PME domain plays a critical role in regulating cell wall rigidity, which is essential for maintaining the integrity of the growing pollen tube. Concurrently, NtLLG4 functions as a key component of the RALF-LLG-ANX-BUPS complex, which acts as a membrane signaling sensor, controlling membrane dynamics and integrity. Together, NtPPME1-NtLLG4 interaction integrates membrane signaling with cell wall rigidity to maintain pollen tube integrity.

## Discussion

In angiosperms, sexual reproduction relies on the structural integrity of the pollen tube, which serves as a tunnel to deliver two sperm cells and undergoes timely rupture to enable double fertilization (*1, 2*). The regulation of cell wall rigidity in pollen tubes is critical, as it must be precisely coordinated with membrane dynamics to maintain both the structural integrity and allow the necessary rupture during fertilization (*1, 2*). Previous studies have well established the key role of the membrane-associated RALF-ANX-BUPS-LLG protein complex and its downstream signaling pathways in governing pollen tube polarity and integrity (*6, 8, 38*). However, the exact mechanisms by which the cell wall cooperates with membrane dynamics to control pollen tube integrity remain largely uncharacterized.

We demonstrated that the pollen-specific protein NtLLG4 functions as a receptor, recognizing NtPPME1 and mediating its unconventional polar exocytosis into the apical apoplast of growing pollen tubes (Fig. 5-7). This interaction is critical for maintaining the polar localization of NtPPME1, which in turn regulates cell wall rigidity, underlying pollen tube growth and integrity. The role of GPI-APs in polar secretion and protein localization has been well-established in various cellular systems, suggesting a conserved mechanism across biological contexts. During pollen tube growth, GPI-APs such as AtLLG2 and AtLLG3 have been identified as chaperones and coreceptors, facilitating the polar secretion of ANX/BUPS to the PM (*38*). This polar targeting is essential for maintaining membrane dynamics and ensuring pollen tube integrity. Similarly, COBL11, another GPI-AP, interacts with the RALF4-ANX-BUPS protein complex to promote the proper localization and polar distribution of its components, further contributing to the stabilization of the pollen tube structure (*40*). These findings underscore the pivotal role of GPI-AP-mediated protein trafficking and localization in regulating pollen tube growth and integrity.

The function of GPI-APs extends beyond pollen tubes, as these proteins are critical for protein localization in other plant tissues (*58, 59*). In synergid cells, LORELEI which is another member of GPI-APs, acts as a chaperone for FERONIA (FER), a homolog of ANX/BUPS, facilitating its trafficking to the PM (*58*). Likewise, in vegetative tissues, LLG1 mediates the transport of FER from the ER to the PM, reinforcing the conserved function of GPI-APs in ensuring the proper localization of essentially functional proteins (*58, 59*). Moreover, GPI-AP-mediated protein trafficking is not unique to plants, while similar pathways have been identified in yeast and mammals (*60*), indicating a broadly conserved mechanism across species. Despite these significant roles, receptor proteins that facilitate the unconventional exocytosis of cargo proteins remain unidentified in animal and yeast systems. The identification of the receptor protein NtLLG4 which mediates the unconventional polar exocytosis of NtPPME1 provide new insights for the investigation and identification of analogous receptor proteins in mammals and yeast, broadening our understanding of this unconventional secretion pathway. Notably, previous research has shown that JIM7-labeled esterified pectin accumulates significantly at the apex of growing pollen tubes in *LLG2/3-RNAi* lines and *cobl11* mutants (*38, 40*). This observation suggests that GPI-APs play an essential role in modulating the composition of the pollen tube tip cell wall, particularly in relation to pectin distribution and rigidity. Given the importance of pectin in maintaining cell wall rigidity and integrity, these findings imply that GPI-APs can regulate membrane dynamics, meanwhile also contribute to the construction and remodeling of the cell wall during pollen tube growth.

Expanding upon these insights, we further reveal that NtLLG4 regulates the polar exocytosis of NtPPME1, a key enzyme involved in regulating the rigidity of pectin during cell wall formation, and contributes to the regulation of the stiffness of pectic cell wall at the pollen tube tip (Fig. 7). Moreover, an optimal level of NtLLG4 is crucial for the accurate targeting and polar exocytosis of NtPPME1 to the pollen tube tip apoplast (Fig. 7). This observation aligns with previous studies demonstrating that the expression level of LLG proteins is a key determinant of pollen tube tip polarity and integrity (*6, 38*). This regulatory mechanism is crucial for governing pollen tube polar growth and cell integrity during fertilization, as it coordinates the interplay between cell wall rigidity and membrane dynamics (Fig. 8). Thus, NtLLG4 emerges as a central molecule that integrates signaling and structural processes required for proper pollen tube growth (*38, 40, 58, 59*). NtLLG4 is involved in the RALF-ANX-BUPS signaling cascade, mediating downstream ROS and Ca²⁺ signaling to sustain membrane integrity (*39*). On the other hand, our findings suggest that it acts as a critical receptor for NtPPME1, facilitating its polar exocytosis and modulating cell wall rigidity. This dual function of NtLLG4 in both signaling and cell wall construction underscores its key role in ensuring proper pollen tube polar growth and fertilization.

Tip-focused rapid pollen tube growth is driven by the accumulation of numerous small apical vesicles, including those involved in exocytic secretion, endocytosis and recycling, all concentrated in the tip region (*18, 45, 61, 62*). Morphological studies using TEM have consistently identified a predominantly homogeneous population of non-coated vesicles with approximately 120 nm in diameter within the pollen tube tip (*23, 29, 43*). A key question that arises is why these various types of apical vesicles, originating from distinct endomembrane trafficking pathways, exhibit such strikingly similar morphologies. We previously identified that the secretion and tip-targeting of NtPPME1 occurs through an unconventional polar exocytosis pathway, in which GDSVs directly mediate the polar exocytosis of NtPPME1 bypassing the TGN in growing pollen tubes (*19*). It is noteworthy that GDSVs exhibit an approximate diameter of 40 nm, as observed in ultrathin-structure images from TEM and 3D tomography of Golgi apparatus (*30, 63*). Therefore, it is highly speculated that GDSVs, despite their smaller size and distinct function, may exhibit unique morphological characteristics that set them apart from other apical vesicles, reflecting their specialized role in mediating the unconventional exocytosis of NtPPME1.

Distinct from conventional TEM sample preparation and imaging approaches, we employed cryo-FIB-SEM imaging combined with 3D tomography and identified two distinct populations of vesicles with contrasting morphologies at the pollen tube tip (Fig. 1). Cryo-FIB-SEM is a powerful technique that allows for the observation of cellular structures in their near-native state by direct sample ultra-thin section and imaging after quick freezing. It greatly preserves their natural morphology and minimizing potential artifacts and morphological changes typically introduced by chemical fixation or embedding resin infiltration (*32–35*). Unlike conventional TEM which often provides two-dimensional images, cryo-FIB-SEM offers three-dimensional high-resolution visualization, allowing for more detailed spatially morphological analysis of complex cellular structures (*33, 64*). The technique achieves continuous sequential sectioning at resolutions as high as 10 nm, facilitating three-dimensional ultrastructural analysis of spatial distribution and organization of organelle and membrane-bound structure morphology across a wide range of biological specimens (*65*). Cryo-FIB-SEM has proven particularly valuable for analyzing vesicle morphology at the pollen tube tip, as demonstrated in our study (Fig. 1). It overcomes the technical challenges of preparing pollen tubes for ultrathin sectioning, where the random spatial distribution of tubes complicates precise sectioning of the tip region. The cryo-FIB-SEM allows for targeted sample preparation, enabling precise identification of the pollen tube tip and the acquisition of continuous serial sections through this critical region (Fig. 1). We utilized advanced cryo-FIB-SEM imaging and 3D tomography to identify and characterize Type II vesicles within the pollen tube tip. These vesicles were found to be ubiquitously distributed and exhibited low electron density and an average diameter of 130 nm (Fig. 1). Our results align with previous studies using conventional TEM, which reported average vesicle diameters of 102 nm, 115 nm, and 128 nm in *Pyrus pyrifolia*, *Papaver rhoeas*, and *Arabidopsis thaliana*, respectively (*23–25*).

More remarkably, our study uncovered a previously undocumented subpopulation of vesicles, designated as Type I vesicles (Fig. 1), and distinguished by their smaller size and high electron density. These vesicles exhibit a remarkable resemblance to GDSVs localized near the Golgi apparatus in the subapical region of pollen tubes (*19, 30, 44*). The rapid sample freezing and afterwards immediate sectioning and imaging by using cryo-FIB-SEM without extra steps of sample substitution and resin embedding minimize the potential risk of morphological changes of endomembrane trafficking vesicles and organelles when compared with conventional chemical fixation or advanced high-pressure freezing and freeze substitution methods (*23, 32, 35, 36, 66*). In addition, cryo-FIB-SEM coupled with 3D tomography generates comprehensive three-dimensional volumes of the pollen tube tip, allowing for the clear observation of different vesicle types and their dynamic behaviors (Fig. 1). Notably, the subcellular localization and distribution pattern of NtPPME1-GFP differs from those of other proteins associated with tip trafficking vesicles, such as SCAMPs and RabA4d (*18, 21, 67, 68*). Unlike these proteins, NtPPME1-GFP fluorescent signals do not prominently accumulate in the tip region of growing pollen tubes (Fig. 2A). This observation, corroborated by multiple previous independent studies (*17, 19, 31, 44*), is likely attributable to two possible reasons: i) the polar exocytosis of NtPPME1 is an exceptionally efficient process characterized by a low incidence of unsuccessful secretion events or recycling within the tip region. Consequently, fluorescent signals from GDSVs carrying NtPPME1-GFP do not accumulate appreciably at the pollen tube tip. This hypothesis is supported by our result demonstrating that disruption of NtPPME1 exocytic secretion markedly increases the fluorescence intensity of a mutated NtPPME1-GFP variant at the pollen tube tip (Fig. 2H panel iv); ii) The polar exocytosis of NtPPME1 and its subsequent release into apoplast, where it plays a critical role in regulating apical cell wall rigidity, occurs as a superfast and transient process. This dynamic behavior is distinct from the trafficking pathways of apical PM-localized or - associated proteins (*18, 21, 67, 68*). Cryo-FIB-SEM provides us a new opportunity to gain unprecedented insights into the populations of apical vesicles in pollen tube tip and greatly enhancing our understanding of the mechanisms underlying pollen tube tip growth and vesicular trafficking.

Polar exocytosis and targeting of proteins, lipids and cell wall components to the pollen tube tip are essential for supporting rapid tube expansion and growth (*8, 19, 69*). In contrast to the conventional exocytic pathway, where proteins are transported from the ER through the Golgi and TGN to the PM, NtPPME1 has been found within GDSVs that bypass the TGN. However, the specific protein trafficking determinants that mediate this unconventional polar exocytic pathway remain largely unknown (*19*). Our study revealed that truncation of either the pro-region or the PME domain of NtPPME1 abolishes its apical localization at the pollen tube tip. Notably, the subcellular localization patterns of the truncated NtPPME1 lacking the pro-region or PME domain differed significantly, suggesting that distinct domains of NtPPME1 may contain specific trafficking signals that guide its trafficking through various intracellular organelles to mediate its unconventional polar exocytosis (Fig. 2). Indeed, this observation is consistent with previous studies (*31*). Furthermore, our systematic analysis identified specific trafficking signals to facilitate the export of NtPPME1 from the ER and Golgi apparatus (Fig. 3). These trafficking determinants are conserved with those previously identified in secretory proteins across eukaryotic cells, suggesting that NtPPME1 follows a conserved protein transport mechanism for secretion through the ER and Golgi (*70, 71*).

In addition, we identified two unique trafficking signals that regulate the unconventional polar transport of NtPPME1 from GDSVs to the pollen tube tip (Fig. 4). In particular, our findings reveal that N-glycosylation plays a crucial role in the unconventional exocytosis of NtPPME1. N-glycosylation is well known for its critical role in the proper localization and function of membrane proteins in the conventional secretory pathway, such as metabotropic glutamate receptors and ion channels (*53, 54, 72, 73*). Our findings expand this role into unconventional trafficking, showing that N-glycosylation of NtPPME1 facilitates its accurate targeting and secretion *via* GDSVs, bypassing the TGN (Fig. 4 and fig. S5C). This adds a novel dimension to our understanding of glycosylation in plant cells, linking it to unconventional polar exocytosis and revealing a broader regulatory mechanism than previously revealed (*54, 74*). Moreover, our finding contrasts with earlier studies focused primarily on conventional secretion pathways, where glycosylation is key for sorting proteins at the TGN (*54, 75*). For instance, glycosylation is crucial for the proper sorting of vacuolar sorting receptors within the TGN in Arabidopsis cells (*54*). However, our results suggest that N-glycosylation plays an equally pivotal role in the function and exocytosis of soluble proteins NtPPME1 that bypass the TGN. This specific role of N-glycosylation highlights its versatility in plant cellular processes and provides a deeper understanding of glycan-mediated protein trafficking.

Taken together, our study reveals a critical mechanism by which GDSV-mediated unconventional polar exocytosis of NtPPME1 coordinates cell wall rigidity with membrane signaling to maintain pollen tube integrity during fertilization. Moreover, we demonstrated that different populations of apical vesicles exist to meet the distinct demands of pollen tube tip growth. These findings advanced our understanding of unconventional polar exocytosis in plant fertilization and provide new insights into the molecular mechanisms governing polar cell growth and morphogenesis. Additionally, elucidating the specific trafficking signals involved in NtPPME1 unconventional exocytosis lays the groundwork for future applications in biotechnology, such as optimizing protein production systems.

## Materials and Methods

### Plant materials and growth conditions

Transgenic tobacco plants expressing NtPPME1-GFP were generated by transforming *Nicotiana tabacum* with the construct *UBQ10_pro_:NtPPME1-GFP*. The primers used for construction of the plasmid are detailed in table S2. *Nicotiana tabacum* plants were grown in a greenhouse at 28°C under a light cycle of 12-h light and 12-h darkness as described before (*76*). *Nicotiana benthamiana* seeds were directly sowed in the soil (Jiffy, EN12580) for germination. Three weeks later, each juvenile plant was transferred to an individual flowerpot and grown in a plant cultivate chamber at 28 °C, 65% humidity, under a light cycle of 16-h light and 8-h darkness with light intensity of 12,000 lux.

### Cryo-FIB-SEM and 3D tomography

Pollen grains were germinated *in vitro* for two hours and vitrified on Au grids (Quantifoil, 200-mesh, gold R2/2) plunge-freezing with Leica EM GP1. Prior to loading the sample, the cryo-stage and cryo anti-contaminator were pre-cooled to -170°C and -190°C, respectively. Subsequently, the specimen grid was mounted into the FIB-SEM (ZEISS Crossbeam 550) cryo-shuttle in the loading station of Quorum PP3010Z cryo-transfer system. This shuttle was then loaded into the pre-chamber of Quorum PP3010Z for sputter coating (5 mA, 60 s) and transferred to the cryo-stage of the ZEISS Crossbeam 550 FIB-SEM. Following transfer, the grid was deposition (30°C,15 s) with platinum precursor using the gas injection system to reduce radiation damage from FIB-SEM. After deposition, pollen tubes on the grid were screened and acquired serial-sections using FIB-SEM. The parameters of SEM imaging was 3.0 kV accelerate voltage, 50 pA electron beam current and secondary electron detector. While 30 kV accelerate voltage, 700 pA ion beam current were used to execute FIB milling and the slice thickness was 30 nm. For model generation, the contours were drawn manually and meshed with the 3dmod program in the IMOD software package (https://bio3d.colorado.edu/imod/doc/guide.html#ParallelProc).

### Transient expression and confocal imaging

Transient expression with growing pollen tubes and tobacco BY-2 protoplasts were carried out essentially as described previously (*77, 78*). Confocal observation and image collection were performed as previously described. The images from pollen tubes or protoplasts expressing fluorescent tagged proteins were collected with a laser level of ≤3% to ensure that the fluorescent signal was within the linear range of detection (typically 1-2% laser power was used). Time-lapse images of growing pollen tubes were collected with minimal time intervals. The primers used for constructing recombinant plasmids were listed in table S2.

### Bioinformatic analysis

Protein sequences were from the NCBI (https://www.ncbi.nlm.nih.gov/). Protein evolutionary conservation analysis was performed using InterPro (https://www.ebi.ac.uk/interpro/). The N-linked glycosylation sites of NtPPME1 were predicted using NetNGlyc (https://services.healthtech.dtu.dk/services/NetNGlyc-1.0/). The phylogenetic tree analysis of AtLLGs and NtLLGs were constructed using MEGA version 7.0. Protein conservation of NtLLGs was analyzed with DNAMAN version 9.0. The 3D structures of NtPPME1 and NtLLG4 were predicted using AlphaFold3 (https://golgi.sandbox.google.com/) and visualized with PyMOL version 2.5 (https://pymol.org/). Prediction results were assessed based on pTM and ipTM values, with the top-ranked model selected for further analysis. Protein-protein interaction interfaces between NtPPME1 and NtLLG4 were predicted using PDBePISA (https://www.ebi.ac.uk/pdbe/pisa/).

### Transcriptome data analysis

The *Nicotiana tabacum* genome (Ntab-TN90) was downloaded from NCBI (https://www.ncbi.nlm.nih.gov/datasets/genome/GCF_000715135.1/) and used as the genome assembly. RNA-seq datasets of *Nicotiana tabacum* (accession numbers: SRR955761, SRR955762, SRR955763, SRR1199069, SRR1199070, SRR1199071, SRR1199072, SRR1199073, SRR1199074, SRR1199121, SRR1199122, SRR1199123, SRR1199124, SRR1199125, SRR1199127, SRR1199128, SRR1199129, SRR1199130, SRR1199132, SRR1199135, SRR1199197, SRR1199198, SRR1199199, SRR1199200, SRR1199202, SRR1199203) were also obtained from NCBI (https://www.ncbi.nlm.nih.gov/sra). The RNA-seq reads were aligned to the Ntab-TN90 reference genome. Gene expression levels were normalized and calculated as fragments per kilobase of transcript per million mapped reads (FPKM) values.

### Transient RNAi plasmid construction

To construct the *NtLLG-RNAi* plasmid under the control of the UBQ promoter, a 258 bp conserved sequence from the *NtLLG4* gene, representing the conserved region of the *NtLLG* family, was amplified in two distinct fragments. These fragments were subsequently cloned into the hairpin RNAi vector pHANNIBAL to form a hairpin RNA structure. The resulting RNAi construct, containing the UBQ promoter and an octopine synthase terminator, was subcloned into the *pBI221-UBQ10_pro_:GFP-NOS* vector to facilitate transient expression.

### Y2H assay

Y2H assays were conducted using the MatchMaker GAL4 Two-Hybrid System 3 (Clontech, 630489) in accordance with the manufacturer’s instructions. The pGBKT7 and pGADT7 vectors were used to carry target cDNAs and then transformed into the *Saccharomyces cerevisiae* strain AH109. After selection on SD –Leu –Trp medium, single transformant colonies were screened for growth on SD –Leu –Trp –His –Ade medium to determine protein interactions. The primers used are listed in table S2.

### LCA

The coding sequence of target genes were amplified and inserted into the pCAMBIA-35S-nLUC and pCAMBIA-35S-cLUC vectors respectively. The plasmids were transformed into *Agrobacterium tumefaciens* GV3101. Transformed bacteria were cultured, harvested and re-suspended in the buffer (10 mM MgCl_2_, 200 μM acetosyringone, 10 mM MES, pH 5.7) to a final concentration of OD_600_ = 0.8-1.0. The suspension was infiltrated into *Nicotiana benthamiana* leaves using a syringe. The tobacco plants were placed in darkness for 24 h and then incubated in a 16 h/8 h light/dark cycle for 24 h. Thereafter, the leaves were sprayed with 1 mM luciferin solution, then kept in darkness for 5 min to quench the fluorescence. A deep cooling CCD imaging apparatus (Tannon, 5200) was used to capture fluorescence images. The primers used for the plasmid construct are listed in table S2.

### FRET assay

For the FRET assay, various constructs, including *UBQ10_pro_:NtPPME1-CFP, UBQ10_pro_:NtLLG4-YFP, UBQ10_pro_:NtLLG4-YFP, UBQ10_pro_:CFP, UBQ10_pro_:YFP, UBQ10_pro_:CFP-YFP* were transiently co-transformed into tobacco BY-2 protoplasts. Following 10-h incubation, FRET analysis was performed using a Leica TCS SP8 confocal system, according to the manufacturer’s instructions, with excitation at 405 nm and 514 nm. The CFP-YFP fusion construct, which includes a short linker between CFP and YFP, served as the positive control, while the co-expression of separate CFP and YFP proteins was used as the negative control. Protoplasts expressing the various CFP and YFP fusion proteins were subjected to photobleaching using a 514 nm laser at full power intensity. FRET efficiency was calculated using the formula FRET_eff_ = (D_post_ – D_pre_)/D_post_, where D_post_ and D_pre_ represent the CFP intensities before and after acceptor bleaching, respectively.

### Protein extraction and HPLC-MS/MS analysis

Germinated genetic transgenic pollen tubes expressing *UBQ10_pro_:NtPPME1-GFP* were collected from ∼200 fresh flowers. After 3-h pollen germination, pollen tubes were frozen in liquid nitrogen, and then resuspended in 2 mL of extraction buffer (50 mM Tris/HCl pH 7.5, 150 mM NaCl, 1 mM MgCl₂, 20% glycerol, 0.2% NP-40, 1× protease inhibitor cocktail). The samples were centrifuged at 13,000 rpm for 30 min and the supernatant was collected. For IP, 50 μL of the supernatant was reserved as input, and the rest was incubated with anti-GFP magnetic beads (Chromotek, gtma) for 2 h at 4°C. The beads were washed twice with extraction buffer and sent for HPLC-MS/MS analysis.

### Co-IP and western blot analysis

The full-length cDNA of *NtPPME1* with its various truncations were tagged with *GFP*, while the full-length of *NtLLG4* was tagged with *Myc*. These chimeric fusion genes were cloned into pCAMBIA expression vectors and transiently co-expressed in *Nicotiana benthamiana* leaves. For protein extraction, 2 g of frozen leaf tissue was ground in liquid nitrogen and homogenized in 2 mL IP buffer containing 50 mM of Tris-HCl, pH 7.4, 150 mM of NaCl, 1 mM of EDTA, 0.5% NP-40 (v:v; Sigma, 127087-87-0), 10% glycerol (v:v) and 1×protease inhibitor cocktail (Roche, 5892970001). The lysates were then incubated with GFP-Trap magnetic agarose beads (ChromoTek, gtma-100) at 4°C for 4 h in a top-to-end rotator. Samples were subsequently analyzed by SDS-PAGE and western blot using anti-GFP (Invitrogen, A11122) and anti-Myc (Sangon Biotech, D110006) antibodies. Chemiluminescence was detected with an image analyzer (Tanon, 5200). The primers used for the plasmid construct are listed in table S2.

### Immunofluorescence labeling

Fixation of *Nicotiana tabacum* pollen tubes, antibody labeling and subsequent immunofluorescence analysis were conducted as previously described (*21, 79*). Briefly, 2-hour germinated pollen tubes, following bombardment, were fixed in a solution containing 3.7% paraformaldehyde, 50 mM Na-phosphate buffer (pH 7.0), 5 mM EGTA, 0.02% Azide. The samples were then incubated overnight at 4°C with JIM5/JIM7 antibodies (1:200, v/v, Abmart, ZW00005/ZW00007) in blocking buffer 2 (0.25% BSA, 0.25% Gelatin, 0.05% NP-40, 0.02% azide in PBS). After washing with blocking buffer 2, the samples were incubated for 1 h with anti-rat lgG antibodies (1:5000, v/v, Life technologies, A-11077). After additional washes, samples were imaged using a Leica TCS SP8 confocal microscope.

### Image processing and analysis

Colocalization analysis between two fluorescent proteins were performed using Fiji software (https://fiji.sc/) with the Pearson-Spearman correlation colocalization plug-in as described previously (*80*). Results were presented either as Pearson correlation coefficients or as Spearman’s rank correlation coefficients, both of which produce r values in the range (-1 to 1), where 0 indicates no discernable correlation and +1 or -1 indicates strong positive or negative correlations, respectively. Signal intensity analysis was measured using plot profile plug-in of Fiji.

### Statistical analysis

Statistical analysis was performed as described in each figure legend. All experiments were performed at least in triplicate (N ≥ 3; exact values indicated in figure legends), with all data points displayed along with the means ± SD. Raw data and statistical analysis for all graphs are presented in table S3.

## Accession numbers

The locus identifiers for the genes mentioned in this article are *NtPPME1* (LOC107768376), *NtLLG1* (LOC107768055), *NtLLG2* (LOC107769788), *NtLLG3* (LOC107800316), *NtLLG4* (LOC107817512), *NtLLG5* (LOC107783567), *NtLLG6* (LOC107783568), *NtLLG7* (LOC107797956), *NtLLG8* (LOC107762859), *NtLLG9* (LOC107814930), *AtLORELEI* (AT4G26466), *AtLLG1* (AT5G56170), *AtLLG2* (AT2G20700), *AtLLG3* (AT4G28280).

## Abbreviations

ANX1/2: ANXUR1/2
BUPS1/2: Buddha’s paper seal ½
cryo-FIB-SEM: cryo-focused ion beam-scanning electron microscope
ER: endoplasmic reticulum
FER: FERONIA
FPKM: fragments per kilobase of transcript per million
FRET: fluorescent resonance energy transfer
GPI-AP: glycosylphosphatidylinositol-anchored protein
GDSVs: Golgi-derived secretory vesicles
HPLC-MS/MS: high performance liquid chromatography-tandem mass spectrometry
IP: immunoprecipitation
KOG: Eukaryotic orthologous groups
LCA: luciferase complementation assay
LLG2/3: LORELEI-LIKE GPI-ANCHORED PROTEIN 2/3
NtLLG4: *Nicotiana tabacum* LORELEI-like-GPI-AP 4
NtPPME1: *Nicotiana tabacum* pollen-specific pectin methylesterase 1
PAE: Predicted aligned error
pLDDT: predicted local distance difference test
PM: plasma membrane
PMEI: PME inhibitor
pTM: predicted template modeling
RC: random coil
RLKs: receptor-like kinases
ROS: reactive oxygen species
SP: signal peptide
TEM: transmission electron microscopy
TGN: *trans*-Golgi network
Y2H: yeast two-hybrid.

## Acknowledgments

We apologize to those whose work could not be cited because of space restrictions. We would like to thank the members of Wang laboratory for stimulating discussions. We also thank Prof. Jiang Tian and Prof. Hong Wu (South China Agricultural University) for providing equipment support. We acknowledge Baolin Qiu and Ruihua Wang (ZEISS Microscopy Customer Center, Guangzhou laboratory) for providing access to the cryo-FIB-SEM facilities.

## Author contributions

Conceptualization: X.W. and Hao Wang. Investigation: X.W., H.W., Y.J., Z.W., C.L., Z.C. and Z.Y. Technical support: Y.J., J.G., L.J., J.H. and L.Z.

Analysis: X.W., H.W., Z.W., Z.Y., J.G. and Hao Wang. Software: X.W. and H.W. Project administration: Hao Wang. Funding acquisition: Hao Wang. Supervision: Hao Wang. Writing-original draft: X.W. Writing-review and editing: X.W. and Hao Wang. All authors approved the final version of the manuscript for submission.

## Data and materials availability

All data needed to evaluate the conclusions in the paper are present in the paper and/or the Supplementary Materials.

## References

1. T. Dresselhaus, S. Sprunck, G. M. Wessel, Fertilization mechanisms in flowering plants. Curr. Biol. 26, 125–139 (2016).

2. J. Zhang, Q. Huang, S. Zhong, A. Bleckmann, J. Huang, X. Guo, Q. Lin, H. Gu, J. Dong, T. Dresselhaus, L. J. Qu, Sperm cells are passive cargo of the pollen tube in plant fertilization. Nat. Plants 3, 17079 (2017).

3. H. J. Li, J. G. Meng, W. C. Yang, Multilayered signaling pathways for pollen tube growth and guidance. Plant Reprod. 31, 31–41 (2018).

4. P. Domingos, P. N. Dias, B. Tavares, M. T. Portes, M. M. Wudick, K. R. Konrad, M. Gilliham, A. Bicho, J. A. Feijó, Molecular and electrophysiological characterization of anion transport in *Arabidopsis thaliana* pollen reveals regulatory roles for pH, Ca^2+^ and GABA. New Phytol. 223, 1353–1371 (2019).

5. Q. Duan, D. Kita, E. A. Johnson, M. Aggarwal, L. Gates, H. M. Wu, A. Y. Cheung, Reactive oxygen species mediate pollen tube rupture to release sperm for fertilization in Arabidopsis. Nat. Commun. 5, 3129 (2014).

6. Z. Ge, Y. Zhao, M. C. Liu, L. Z. Zhou, L. Wang, S. Zhong, S. Hou, J. Jiang, T. Liu, Q. Huang, J. Xiao, H. Gu, H. M. Wu, J. Dong, T. Dresselhaus, A. Y. Cheung, L. J. Qu, LLG2/3 are co-receptors in BUPS/ANX-RALF signaling to regulate Arabidopsis pollen tube integrity. Curr. Biol. 29, 3256–3265 (2019).

7. J. Sekereš, P. Pejchar, J. Šantrůček, N. Vukašinović, V. Žárský, M. Potocký, Analysis of exocyst subunit EXO70 family reveals distinct membrane polar domains in tobacco pollen tubes. Plant Physiol. 173, 1659–1675 (2017).

8. Z. Ge, T. Bergonci, Y. Zhao, Y. Zou, S. Du, M. C. Liu, X. Luo, H. Ruan, L. E. García-Valencia, S. Zhong, S. Hou, Q. Huang, L. Lai, D. S. Moura, H. Gu, J. Dong, H. M. Wu, T. Dresselhaus, J. Xiao, A. Y. Cheung, L. J. Qu, Arabidopsis pollen tube integrity and sperm release are regulated by RALF-mediated signaling. Science 358, 1596–1600 (2017).

9. M. A. Mecchia, G. Santos-Fernandez, N. N. Duss, S. C. Somoza, A. Boisson-Dernier, V. Gagliardini, A. Martínez-Bernardini, T. N. Fabrice, C. Ringli, J. P. Muschietti, U. Grossniklaus, RALF4/19 peptides interact with LRX proteins to control pollen tube growth in Arabidopsis. Science 358, 1600–1603 (2017).

10. Z. Zhou, H. Shi, B. Chen, R. Zhang, S. Huang, Y. Fu, Arabidopsis RIC1 severs actin filaments at the apex to regulate pollen tube growth. Plant Cell 27, 1140–1161 (2015).

11. Y. J. Xu, T. Luo, P. M. Zhou, W. Q. Wang, W. C. Yang, H. J. Li, Pollen-expressed RLCKs control pollen tube burst. Plant Commun 5, 100934 (2024).

12. M. Lenartowska, M. I. Rodríguez-García, E. Bednarska, Immunocytochemical localization of esterified and unesterified pectins in unpollinated and pollinated styles of *Petunia hybrida Hort*. Planta 213, 182–191 (2001).

13. Y. Chebli, M. Kaneda, R. Zerzour, A. Geitmann, The cell wall of the Arabidopsis pollen tube-spatial distribution, recycling, and network formation of polysaccharides. Plant Physiol. 160, 1940–1955 (2012).

14. J. C. Mollet, C. Leroux, F. Dardelle, A. Lehner, Cell wall composition, biosynthesis and remodeling during pollen tube growth. Plants (Basel) 2, 107–147 (2013).

15. M. Cascallares, N. Setzes, F. Marchetti, G. A. López, A. M. Distéfano, M. Cainzos, E. Zabaleta, G. C. Pagnussat, A complex journey: cell wall remodeling, interactions, and integrity during pollen tube growth. Front. Plant Sci. 11, 599247 (2020).

16. P. B. Adhikari, X. Liu, R. D. Kasahara, Mechanics of pollen tube elongation: A perspective. Front. Plant Sci. 11, 589712 (2020).

17. N. Stroppa, E. Onelli, P. Moreau, L. Maneta-Peyret, V. Berno, E. Cammarota, R. Ambrosini, M. Caccianiga, M. Scali, A. Moscatelli, Sterols and sphingolipids as new players in cell wall building and apical growth of *Nicotiana tabacum L.* pollen tubes. Plants (Basel) 12, 8 (2022).

18. X. Weng, Y. Shen, L. Jiang, L. Zhao, H. Wang, Spatiotemporal organization and correlation of tip-focused exocytosis and endocytosis in regulating pollen tube tip growth. Plant Sci. 330, 111633 (2023).

19. H. Wang, X. Zhuang, X. Wang, A. H. Law, T. Zhao, S. Du, M. M. Loy, L. Jiang, A distinct pathway for polar exocytosis in plant cell wall formation. Plant Physiol. 172, 1003–1018 (2016).

20. L. Zonia, T. Munnik, Vesicle trafficking dynamics and visualization of zones of exocytosis and endocytosis in tobacco pollen tubes. *J*, Exp. Bot. 59, 861–873 (2008).

21. H. Wang, Y. C. Tse, A. H. Law, S. S. Sun, Y. B. Sun, Z. F. Xu, S. Hillmer, D. G. Robinson, L. Jiang, Vacuolar sorting receptors (VSRs) and secretory carrier membrane proteins (SCAMPs) are essential for pollen tube growth. Plant J. 61, 826–838 (2010).

22. Y. Cai, X. Zhuang, J. Wang, H. Wang, S. K. Lam, C. Gao, X. Wang, L. Jiang, Vacuolar degradation of two integral plasma membrane proteins, AtLRR84A and OsSCAMP1, is cargo ubiquitination-independent and prevacuolar compartment-mediated in plant cells. Traffic 13, 1023–1040 (2012).

23. P. F. Jia, H. J. Li, W. C. Yang, Transmission electron microscopy (TEM) to study histology of pollen and pollen tubes. Methods Mol. Biol. 1669, 181–189 (2017).

24. A. Geitmann, V. E. Franklin-Tong, A. C. Emons, The self-incompatibility response in *Papaver rhoeas* pollen causes early and striking alterations to organelles. Cell Death Differ. 11, 812–822 (2004).

25. C. Shi, D. Wang, Y. Guan, H. Qu, Dissection and ultramicroscopic observation of an apical pollen tube of *Pyrus*. Plant Reprod. 35, 1–8 (2022).

26. 26. R. Malhó, P. C. Coelho, E. Pierson, J. Derksen, “Endocytosis and membrane recycling in pollen tubes” in Plant Endocytosis, J. Šamaj, F. Baluška, D. Menzel, Eds. (Springer Berlin Heidelberg, Berlin, Heidelberg, 2006), pp. 277–291.

27. T. Ketelaar, M. E. Galway, B. M. Mulder, A. M. Emons, Rates of exocytosis and endocytosis in Arabidopsis root hairs and pollen tubes. J. Microsc. 231, 265–273 (2008).

28. J. Derksen, T. Rutten, I. K. Lichtscheidl, A. H. N. de Win, E. S. Pierson, G. Rongen, Quantitative analysis of the distribution of organelles in tobacco pollen tubes: implications for exocytosis and endocytosis. Protoplasma 188, 267–276 (1995).

29. F. Liao, L. Wang, L. B. Yang, X. Peng, M. Sun, NtGNL1 plays an essential role in pollen tube tip growth and orientation likely *via* regulation of post-Golgi trafficking. PLoS One 5, e13401 (2010).

30. B. H. Kang, E. Nielsen, M. L. Preuss, D. Mastronarde, L. A. Staehelin, Electron tomography of RabA4b- and PI-4Kβ1-labeled *trans* Golgi network compartments in Arabidopsis. Traffic 12, 313–329 (2011).

31. M. Bosch, A. Y. Cheung, P. K. Hepler, Pectin methylesterase, a regulator of pollen tube growth. Plant Physiol. 138, 1334–1346 (2005).

32. N. Vidavsky, A. Akiva, I. Kaplan-Ashiri, K. Rechav, L. Addadi, S. Weiner, A. Schertel, Cryo-FIB-SEM serial milling and block face imaging: Large volume structural analysis of biological tissues preserved close to their native state. J. Struct. Biol. 196, 487–495 (2016).

33. D. Spehner, A. M. Steyer, L. Bertinetti, I. Orlov, L. Benoit, K. Pernet-Gallay, A. Schertel, P. Schultz, Cryo-FIB-SEM as a promising tool for localizing proteins in 3D. J. Struct. Biol. 211, 107528 (2020).

34. O. Klykov, D. Bobe, M. Paraan, J. D. Johnston, C. S. Potter, B. Carragher, M. Kopylov, A. J. Noble, *In situ* cryo-FIB/SEM specimen preparation using the waffle method. Bio Protoc. 12, e4544 (2022).

35. A. Schertel, N. Snaidero, H. M. Han, T. Ruhwedel, M. Laue, M. Grabenbauer, W. Möbius, Cryo FIB-SEM: volume imaging of cellular ultrastructure in native frozen specimens. J. Struct. Biol. 184, 355–360 (2013).

36. A. Bullen, R. R. Taylor, B. Kachar, C. Moores, R. A. Fleck, A. Forge, Inner ear tissue preservation by rapid freezing: improving fixation by high-pressure freezing and hybrid methods. Hear Res 315, 49–60 (2014).

37. S. Liang, M. L. Hu, H. C. Lin, H. J. He, X. P. Ning, P. P. Peng, G. H. Lu, S. L. Sun, X. J. Wang, Y. Q. Wang, H. Wu, Transcriptional regulations of pollen tube reception are associated with the fertility of the ginger species *Zingiber zerumbet* and *Zingiber corallinum*. Front. Plant Sci. 14, 1099250 (2023).

38. H. Feng, C. Liu, R. Fu, M. Zhang, H. Li, L. Shen, Q. Wei, X. Sun, L. Xu, B. Ni, C. Li, LORELEI-LIKE GPI-ANCHORED PROTEINS 2/3 regulate pollen tube growth as chaperones and coreceptors for ANXUR/BUPS receptor kinases in Arabidopsis. Mol. Plant 12, 1612–1623 (2019).

39. Q. Gao, C. Wang, Y. Xi, Q. Shao, C. Hou, L. Li, S. Luan, RALF signaling pathway activates MLO calcium channels to maintain pollen tube integrity. Cell Res. 33, 71–79 (2023).

40. H. Li, Y. Yang, H. Zhang, C. Li, P. Du, M. Bi, T. Chen, D. Qian, Y. Niu, H. Ren, L. An, Y. Xiang, The Arabidopsis GPI-anchored protein COBL11 is necessary for regulating pollen tube integrity. Cell Rep. 42, 113353 (2023).

41. P. K. Hepler, C. M. Rounds, L. J. Winship, Control of cell wall extensibility during pollen tube growth. Mol. Plant 6, 998–1017 (2013).

42. L. Zonia, T. Munnik, Understanding pollen tube growth: the hydrodynamic model versus the cell wall model. Trends Plant Sci. 16, 347–352 (2011).

43. T. Ndinyanka Fabrice, A. Kaech, G. Barmettler, C. Eichenberger, J. P. Knox, U. Grossniklaus, C. Ringli, Efficient preparation of Arabidopsis pollen tubes for ultrastructural analysis using chemical and cryo-fixation. BMC Plant Biol. 17, 176 (2017).

44. H. Wang, X. Zhuang, Y. Cai, A. Y. Cheung, L. Jiang, Apical F-actin-regulated exocytic targeting of NtPPME1 is essential for construction and rigidity of the pollen tube cell wall. Plant J. 76, 367–379 (2013).

45. X. Weng, H. Wang, Apical vesicles: Social networking at the pollen tube tip. Reprod. Breed. 2, 119–124 (2022).

46. J. Pelloux, C. Rustérucci, E. J. Mellerowicz, New insights into pectin methylesterase structure and function. Trends Plant Sci. 12, 267–277 (2007).

47. R. P. Jolie, T. Duvetter, A. M. Van Loey, M. E. Hendrickx, Pectin methylesterase and its proteinaceous inhibitor: a review. Carbohydr Res 345, 2583–2595 (2010).

48. Y. Wang, D. Zhang, L. Huang, Z. Zhang, Q. Shi, J. Hu, G. He, X. Guo, H. Shi, L. Liang, Uncovering the interactions between PME and PMEI at the gene and protein levels: Implications for the design of specific PMEI. J. Mol. Model. 29, 286 (2023).

49. S. Wolf, T. Rausch, S. Greiner, The N-terminal pro region mediates retention of unprocessed type-I PME in the Golgi apparatus. Plant J. 58, 361–375 (2009).

50. S. L. Hanton, L. Renna, L. E. Bortolotti, L. Chatre, G. Stefano, F. Brandizzi, Diacidic motifs influence the export of transmembrane proteins from the endoplasmic reticulum in plant cells. Plant Cell 17, 3081–3093 (2005).

51. O. Nufer, F. Kappeler, S. Guldbrandsen, H. P. Hauri, ER export of ERGIC-53 is controlled by cooperation of targeting determinants in all three of its domains. J. Cell Sci. 116, 4429–4440 (2003).

52. K. Sato, A. Nakano, Emp47p and its close homolog Emp46p have a tyrosine-containing endoplasmic reticulum exit signal and function in glycoprotein secretion in *Saccharomyces cerevisiae*. Mol. Biol. Cell 13, 2518–2532 (2002).

53. J. A. Verdinez, J. A. Sebag, Role of N-linked glycosylation in PKR2 trafficking and signaling. Front. Neurosci. 15, 730417 (2021).

54. J. Shen, Y. Ding, C. Gao, E. Rojo, L. Jiang, N-linked glycosylation of AtVSR1 is important for vacuolar protein sorting in Arabidopsis. Plant J. 80, 977–992 (2014).

55. E. Pascual-Morales, P. Jiménez-Chávez, J. E. Olivares-Grajales, L. Sarmiento-López, W. R. García-Niño, A. López-López, P. H. Goodwin, J. Palacios-Martínez, A. I. Chávez-Martínez, L. Cárdenas, Role of a LORELEI-like gene from *Phaseolus vulgaris* during a mutualistic interaction with *Rhizobium tropici*. PLoS One 18, e0294334 (2023).

56. S. A. Lancelle, P. K. Hepler, Ultrastructure of freeze-substituted pollen tubes of *Lilium longiflorum*. Protoplasma 167, 215–230 (1992).

57. T. Ishii, T. Matsunaga, Pectic polysaccharide rhamnogalacturonan II is covalently linked to homogalacturonan. Phytochemistry 57, 969–974 (2001).

58. C. Li, F. L. Yeh, A. Y. Cheung, Q. Duan, D. Kita, M. C. Liu, J. Maman, E. J. Luu, B. W. Wu, L. Gates, M. Jalal, A. Kwong, H. Carpenter, H. M. Wu, Glycosylphosphatidylinositol-anchored proteins as chaperones and co-receptors for FERONIA receptor kinase signaling in Arabidopsis. Elife 4, e06587 (2015).

59. X. Liu, C. Castro, Y. Wang, J. Noble, N. Ponvert, M. Bundy, C. Hoel, E. Shpak, R. Palanivelu, The role of LORELEI in pollen tube reception at the iInterface of the synergid cell and pollen tube requires the modified eight-cysteine motif and the receptor-like kinase FERONIA. Plant Cell 28, 1035–1052 (2016).

60. T. Kinoshita, M. Fujita, Biosynthesis of GPI-anchored proteins: special emphasis on GPI lipid remodeling. J Lipid Res 57, 6–24 (2016).

61. L. Zhao, M. S. Rehmani, H. Wang, Exocytosis and endocytosis: Yin-yang crosstalk for sculpting a dynamic growing pollen tube tip. Front. Plant Sci. 11, 572848 (2020).

62. J. Guo, Z. Yang, Exocytosis and endocytosis: coordinating and fine-tuning the polar tip growth domain in pollen tubes. *J*, Exp. Bot. 71, 2428–2438 (2020).

63. Q. Rui, X. Tan, F. Liu, Y. Li, X. Liu, B. Li, J. Wang, H. Yang, L. Qiao, T. Li, S. Fang, R. Gao, W. Wang, S. Y. Bednarek, Y. Bao, Syntaxin of plants31 (SYP31) and SYP32 is essential for Golgi morphology maintenance and pollen development. Plant Physiol. 186, 330–343 (2021).

64. N. Scher, K. Rechav, P. Paul-Gilloteaux, O. Avinoam, *In situ* fiducial markers for 3D correlative cryo-fluorescence and FIB-SEM imaging. iScience 24, 102714 (2021).

65. V. Baena, R. Conrad, P. Friday, E. Fitzgerald, T. Kim, J. Bernbaum, H. Berensmann, A. Harned, K. Nagashima, K. Narayan, FIB-SEM as a volume electron microscopy approach to study cellular architectures in SARS-CoV-2 and other viral infections: A practical primer for a virologist. Viruses 13, 611 (2021).

66. E. Kellenberger, The potential of cryofixation and freeze substitution: observations and theoretical considerations. J. Microsc. 161, 183–203 (1991).

67. Y. J. Lee, A. Szumlanski, E. Nielsen, Z. Yang, Rho-GTPase-dependent filamentous actin dynamics coordinate vesicle targeting and exocytosis during tip growth. J. Cell Biol. 181, 1155–1168 (2008).

68. A. L. Szumlanski, E. Nielsen, The Rab GTPase RabA4d regulates pollen tube tip growth in Arabidopsis thaliana. Plant Cell 21, 526–544 (2009).

69. F. Hempel, I. Stenzel, M. Heilmann, P. Krishnamoorthy, W. Menzel, R. Golbik, S. Helm, D. Dobritzsch, S. Baginsky, J. Lee, W. Hoehenwarter, I. Heilmann, MAPKs influence pollen tube growth by controlling the formation of phosphatidylinositol 4,5-bisphosphate in an apical plasma membrane domain. Plant Cell 29, 3030–3050 (2017).

70. C. K. Barlowe, E. A. Miller, Secretory protein biogenesis and traffic in the early secretory pathway. Genetics 193, 383–410 (2013).

71. N. Gomez-Navarro, E. Miller, Protein sorting at the ER-Golgi interface. J. Cell Biol. 215, 769–778 (2016).

72. D. H. Park, S. Park, J. M. Song, M. Kang, S. Lee, M. Horak, Y. H. Suh, N-linked glycosylation of the mGlu7 receptor regulates the forward trafficking and transsynaptic interaction with Elfn1. FASEB J. 34, 14977–14996 (2020).

73. J. V. Li, C. A. Ng, D. Cheng, Z. Zhou, M. Yao, Y. Guo, Z. Y. Yu, Y. Ramaswamy, L. A. Ju, P. W. Kuchel, M. P. Feneley, D. Fatkin, C. D. Cox, Modified N-linked glycosylation status predicts trafficking defective human Piezo1 channel mutations. *Commun*. Biol. 4, 1038 (2021).

74. R. Strasser, G. Seifert, M. S. Doblin, K. L. Johnson, C. Ruprecht, F. Pfrengle, A. Bacic, J. M. Estevez, Cracking the “Sugar Code”: A snapshot of N-and O-glycosylation pathways and functions in plants cells. Front. Plant Sci. 12, 640919 (2021).

75. Y. Guo, D. W. Sirkis, R. Schekman, Protein sorting at the *trans*-Golgi network. Annu Rev Cell Dev Biol 30, 169–206 (2014).

76. H. Yan, M. Zhuang, X. Xu, S. Li, M. Yang, N. Li, X. Du, K. Hu, X. Peng, W. Huang, H. Wu, Y. C. Tse, L. Zhao, H. Wang, Autophagy and its mediated mitochondrial quality control maintain pollen tube growth and male fertility in Arabidopsis. Autophagy 19, 768–783 (2023).

77. H. Wang, L. Jiang, Transient expression and analysis of fluorescent reporter proteins in plant pollen tubes. Nat. Protoc. 6, 419–426 (2011).

78. Y. Miao, L. Jiang, Transient expression of fluorescent fusion proteins in protoplasts of suspension cultured cells. Nat. Protoc. 2, 2348–2353 (2007).

79. H. Wang, X. H. Zhuang, S. Hillmer, D. G. Robinson, L. W. Jiang, Vacuolar sorting receptor (VSR) proteins reach the plasma membrane in germinating pollen tubes. Mol. Plant 4, 845–853 (2011).

80. A. P. French, S. Mills, R. Swarup, M. J. Bennett, T. P. Pridmore, Colocalization of fluorescent markers in confocal microscope images of plant cells. Nat. Protoc. 3, 619–628 (2008).

